# An *Orobanche cernua* x *Orobanche cumana* segregating population provides insight into the regulation of germination specificity in a parasitic plant

**DOI:** 10.1101/2022.03.22.485355

**Authors:** Hailey Larose, Dina Plakhine, Nathan Wycoff, Na Zhang, Caitlin Conn, David C. Nelson, Hanan Eizenberg, Daniel M. Joel, Yaakov Tadmor, James H. Westwood

## Abstract

Germination of seeds of *Orobanche* species requires specific chemicals exuded by host roots. A family of “divergent” *KARRIKIN INSENSITIVE2* (*KAI2d*) genes encode proteins that recognize strigolactone (SL) class germination simulants. We explored specificity of germination stimulant detection by analyzing interspecific segregants of a cross between *Orobanche cernua* and *O. cumana*, closely related species that differ in stimulant response. *O. cernua* parasitizes tomato and germinates in response to the SL orobanchol, while *O. cumana* parasitizes sunflower and responds to dehydrocostus lactone (DCL). *KAI2d* genes were catalogued in parents and in segregants that showed stimulant specificity. *KAI2d* genes were also functionally assayed in the *Arabidopsis kai2* mutant background. We identified five full-length *KAI2d* genes in *O. cernua* and eight in *O. cumana*. The *O. cernua KAI2d2*, as well as its ortholog in *O. cumana*, are associated with SL perception. A cluster of *O. cumana KAI2d* genes was genetically linked to DCL perception, although no specific receptor gene was identified by heterologous complementation. These findings support the *KAI2d*-mediated perception of SLs, but fall short of explaining how *O. cumana* perceives DCL. The ability of some *O. cumana KAI2d* genes to detect SLs points to the involvement of additional factors in regulating stimulant specificity.

## Introduction

Parasitic plants cause major damage to agriculture in many areas of the world (Parker, 2013). This damage results in annual losses of approximately 1 billion US dollars, and affects more than a hundred million people (Parker, 2009; Hegenauer *et al*., 2017). The most economically destructive parasitic weeds belong to the family Orobanchaceae, which includes the genera *Striga, Phelipanche and Orobanche* (Parker, 2012). The impact of these weeds can be attributed to multiple factors including their underground location on the host root where they grow undetected while disproportionately limiting host growth (Ishida *et al*., 2016; Spallek *et al*., 2017), and their ability to produce hundreds of thousands of tiny seeds that can persist for years in the soil. Ultimately, they remain a constraint on agriculture because there is currently a lack of effective, affordable strategies to control these parasites. One promising, but yet unrealized, approach to parasitic weed control is to disrupt the host-parasite communication that regulates parasite seed germination (e.g., Samejima *et al*., 2016; Waters, 2017).

Obligate parasitic weeds of the *Orobanchaceae* have evolved to ensure that their seeds germinate only in the presence of a host root. The first requirement for *Orobanche* seed germination is a process termed conditioning, which is a several-day period of exposure to appropriate temperature and moisture, and includes seed imbibition that results in seed swelling and opening of the micropyle (Joel *et al*., 2011a). Once conditioning is complete, seeds are ready to perceive a host-derived chemical that will trigger germination. For most weedy members of *Orobanchaceae* this germination stimulant is a strigolactone (SL). Parasitic plants have evolved a mechanism(s) to detect specific SL variants, singly or in combinations (Yoneyama *et al*., 2013). This signal perception is crucial, because once germinated, the parasite radicle has only a few days to contact the host-root and form a haustorial connection before the seedling’s limited nutrients are exhausted (Westwood *et al*., 2010).

Much work has focused on understanding the mechanism by which parasites are able to detect SLs in their environment. Briefly, key SL receptors for parasite germination are α/β-hydrolases, members of a gene family called *KAI2* (*KARRIKIN INSENSITIVE 2*; also termed *HTL, HYPOSENSITIVE TO LIGHT*) (Conn *et al*., 2015; Toh *et al*., 2015; Tsuchiya *et al*., 2015; Yao *et al*., 2017). *KAI2* genes are evolutionarily related to *D14 (DWARF14*) (Bythell-Douglas *et al*., 2017), which functions in the perception of endogenous SLs to regulate shoot branching and other developmental and environmental responses in angiosperms (Umehara *et al*., 2008; Koltai, 2014; Yang *et al*., 2019). KAI2 appears to be adapted to detect other molecules, including external chemical signals such as karrikins, which are found in smoke (Waters & Smith, 2013; De Cuyper *et al*., 2017). While the specific details of ligand-receptor binding and signaling are still being worked out, a consensus model for KAI2/D14 function is clear (Burger & Chory, 2020; Machin *et al*., 2020). The receptor protein binds the SL with the D ring oriented deep in the binding pocket, resulting in hydrolysis of the D ring. At some point the receptor recruits an F-box protein (MAX2, MORE AXILLARY GROWTH 2) and the complex associates with a protein of the SMAX1-LIKE (SMXL) family, leading to the ubiquitination and subsequent degradation of the SMXL (Khosla *et al*., 2020; Wang *et al*., 2020). Based on genetic studies in *Arabidopsis*, the KAI2 mediated degradation of SMAX1 (SUPPRESSOR OF MAX2 1) ultimately leads to germination (Stanga *et al*., 2013; Bunsick *et al*., 2020).

The ability of Orobanchaceae parasite KAI2/HTL receptors to function in host detection was enabled by an expansion of the gene family and evolution of the receptors themselves to recognize host-derived SLs (Conn *et al*., 2015). *KAI2* paralogs from these parasitic species have been classified into three major phylogenic clades based upon their rates of evolutionary selection: conserved (*KAI2c*), intermediate (*KAI2i*) and divergent (*KAI2d*) (Conn *et al*., 2015). Limited sampling of *KAI2* genes in parasitic plants and assays in transgenic gene complementation studies suggest the three *KAI2* clades have different ligand specificities (Conn *et al*., 2015; Toh *et al*., 2015). *KAI2c* proteins have been proposed to respond to a yet unknown KAI2 ligand endogenous to plants, *KAI2i* respond to karrikins, and the fastest-evolving, parasit-especific *KAI2d* proteins respond to SLs (Tsuchiya *et al*., 2015; Conn & Nelson, 2016). Duplication of *KAI2d* genes in parasitic plants and specialization for specific SL stimulants from different hosts provides an explanation for how parasite species could evolve new host specificities. For example, *S. hermonthica* has at least 13 *KAI2d* genes (Conn *et al*., 2015), with multiple versions (*ShHTL4-9*) able to respond to SL stimulants *in vitro* (Tsuchiya *et al*., 2015) and in *Arabidopsis kai2* complementation assays, but *ShHTL7* appears to confer the highest SL sensitivity to *Arabidopsis* (Toh *et al*., 2015). *Phelipanche ramosa* has five *KAI2d* genes, two of which respond to SLs in heterologous assays (de Saint Germain *et al*., 2021).

Research on stimulant perception is limited by a lack of tools for genetic analyses and manipulation (Nelson, 2021). There are no inbred lines differing in stimulant specificity that could support traditional genetic identification of important loci. The lack of reliable parasite transformation protocols needed for gain- or loss-of-function studies necessitates that all *in vivo* assays of parasite KAI2d specificity and function be conducted in non-parasite (*Arabidopsis*) systems. Conclusions from such studies must be viewed with caution, as a KAI2d protein may function differently in *Arabidopsis* than in its native cellular environment.

To address this gap and evaluate germination specificity in a parasitic plant genetic system, we used the *Orobanche cernua* / *O. cumana* complex. These species present a valuable contrast in that they are closely related, having previously been considered two forms of the same species (Parker & Riches, 1993). *Orobanche cernua* parasitizes Solanaceous crops and germinates in response to SLs such as orobanchol (Oro), a major SL exudate from tomato (*Solanum lycopersicum*) (Dor *et al*., 2011). In contrast, *O. cumana* parasitizes sunflower (*Helianthus annuus*) and germinates in response to dehydrocostus lactone (DCL) rather than natural SL stimulants (Joel *et al*., 2011b). Both species respond to the synthetic strigolactone racemate, *rac*-GR24 (Plakhine *et al*., 2012). These species can be crossed to produce fertile offspring that segregate for stimulant response (Plakhine *et al*., 2012). From this study, F_3_ families were selected based on their germination responses to DCL, the natural SL Orobanchol (Oro), and the synthetic SL analog, *rac*-GR24.

Here we describe experiments using segregants derived from the cross *O. cernua* x *O. cumana* to identify the key components underlying germination specificity. We hypothesized that some KAI2d proteins may function as receptors for Oro, while others (i.e., from *O. cumana*) would be specific for DCL. We sequenced transcriptomes of both species and found differences in the numbers of *KAI2d* genes. We genotyped segregants differing in stimulant response phenotype and identified *KAI2d* genes that were correlated with stimulant response. Finally, we cloned and expressed each *KAI2* gene from both species in a heterologous system, identifying multiple forms that recognize SLs, but no receptor for DCL. Taken together, our results indicate that *KAI2d* proteins are involved in perception of SLs in parasitic plants, but suggest other layers of regulation are involved in conferring specificity to DCL.

## Materials and Methods

### Seed sources

*Orobanche cernua* seeds were collected in tomato fields in Israel in 1994 and *O. cumana* seeds were collected in sunflower fields in Israel in 1997. *Orobanche cernua* and *O. cumana* were then multiplied each year in a net-house at the Newe Ya’ar Research Center on tomato and sunflower, respectively.

### Tissue collection for RNA-sequencing

To obtain sequences of genes expressed during conditioning and germination, we sequenced transcriptomes of *O. cernua* and *O. cumana* seeds at three time points spanning the perception of the germination signal: during conditioning (“conditioning”), at completion of conditioning (“fully conditioned”), and following exposure to germination stimulants (“stimulated”). The final stage was further divided into treatments with either a species-specific stimulant DCL (Sigma-Aldrich) or Oro (Initially from Koichi Yoneyama, then from StrigoLab), or a racemic mixture of (+) and (-)- GR24 (*rac*-GR24), which is a SL analog used as a universal stimulant for studies of *Orobanche, Phelipanche* and *Striga* (Wigchert *et al*., 1999; Yoneyama *et al*., 2010). Seeds *O. cernua* and *O. cumana* from parental lines were surface sterilized according to Plakhine *et al*. (2012). For the “conditioning” treatment, seeds were placed in a Petri dish as described and stored in darkness at 23°C for 1, 3 or 5 days, and seeds from each time point were pooled. For “fully conditioned”, seeds were treated as for conditioning, but held for 7 days. For “stimulated”, conditioned seeds were exposed for 4 or 8 hrs to either DCL (10^-7^ μM), Oro (10^-8^ μM), or *rac*-GR24 (10^-6^ μM), where concentrations were previously determined to yield optimal germination. In all cases seeds were pooled to comprise two biological replicates. RNA was extracted with a Spectrum Plant Total RNA kit (Sigma-Aldrich). Strand-specific RNA-seq libraries were constructed, barcoded and sequenced to obtain 100bp paired-end (PE) reads on two lanes of Illumina HiSeq 2000 sequencing system at the Technion Genome Center in Haifa, Israel.

### *De novo* transcriptome assembly of *O. cernua* and *O. cumana*

Raw read quality was assessed using FastQC (Andrews, 2010). Prior to assembly, raw reads were trimmed using Trimmomatic (Bolger *et al*., 2014). Reads of at least 50bp that retained their PE mate were subjected to *de novo* transcriptome assembly using the Trinity software package (version 2.4.0) with default parameters (Haas *et al*., 2013). Transcriptomes were assembled for all sequenced stages of each species. Raw reads were mapped back to the *de novo* transcriptomes using Bowtie2 software and had a mapping efficiency greater than ninety percent for both species (Langmead & Salzberg, 2012). The transcriptomes of *O. cernua* and *O. cumana* each produced over 200,000 contigs (Supporting Information Table S1) and were roughly equivalent in sequencing depth and number of predicted ESTs, with 103,570 ESTs identified for *O. cernua* and 110,019 for *O. cumana*. The Core Eukaryotic Genes Mapping Approach (CEGMA) pipeline was used to estimate the completeness of transcriptome assemblies (Parra *et al*., 2007), and each transcriptome assembled 99% of CEGMA proteins, suggesting high transcriptome completeness.

### *KAI2* gene identification

*KAI2* and *D14* gene sequences were identified for each species through tblastx searches using *Arabidopsis thaliana KAI2* sequences with an e-value of < 10^-5^ (Altschul *et al*., 1997). All contig hits were extracted from each transcriptome and aligned against the *O. cernua* and *O. cumana KAI2d* genes identified by Conn, *et al*. (2015). Sequences were validated by Reverse Transcriptase (RT) PCR amplification using RNA isolated from seven-day conditioned *O. cernua* and *O. cumana*, and primers described in Supporting Information Table S2, followed by Sanger sequencing.

### Phylogenetic reconstruction

In addition to the *KAI2* sequences from *O. cernua* and *O. cumana* reported in this study, predicted coding sequences of *KAI2* and *D14* from species in the Orobanchaceae were collected from Conn *et al*. (2015) and Yoshida *et al*. (2019). *Orobanche cernua KAI2d6* was removed from the alignment as it is greatly reduced length and likely a pseudogene. Sequences were manually aligned with respect to the predicted amino acid sequences and ends trimmed to minimize regions of unclear homology. Relative to *Arabidopsis thaliana KAI2*, codons 7 – 266 were retained. The final alignment contained 80 *KAI2* and 11 *D14* sequences and comprised 801 characters.

MrBayes (version 3.2; Ronquist *et al*., 2012) was used to reconstruct a Bayesian phylogeny of aligned *KAI2* and *D14* sequences. Two independent MCMC runs were used with the General Time Reversible + Γ + I model of evolution for 2.0 x 10^7^ generations, with sampling every 1.0 x 10^4^ generations. The first 25% of trees were discarded as burnin, and the remaining trees were used to create a consensus tree. The consensus tree was rerooted with the monophyletic *D14* clade as outgroup.

### Segregating population

Crosses of *O. cernua* and *O. cumana* are described in Plakhine *et al*. (2012), and F_2_ of this cross were self-pollinated to produce the F_3_ families, at which point the stimulant response phenotype segregates. The segregation seen between and within the F_3_ generation is attributed to expression of the stimulant perception mechanism in the maternally derived perisperm tissue of the seeds (Plakhine *et al*., 2012). For this reason, genotyping was conducted on the F_2_ segregants while phenotyping was conducted on the F_3_ families. A total of 94 F_2_ segregants were evaluated.

### Genotyping *O. cernua* x *O. cumana* F_2_ plants

Floral shoot tissue from *O. cernua* and *O. cumana* parental and F_2_ segregant plants was collected and genomic DNA extracted using a cetyltrimethylammonium bromide (CTAB)-based extraction method (Doyle & Doyle, 1987). Tissue was ground under liquid nitrogen using a mortar and pestle and mixed with 500μL of 2X CTAB buffer. Purified DNA was resuspended with 1X Tris-EDTA buffer (10mM Tris-HCL, 0.1mM EDTA, pH 8.0) overnight at 4 °C. The quality and concentration of the DNA were checked using Nanodrop One (ThermoFisher) and gel electrophoresis.

Due to the large number of segregants potentially harboring several *KAI2d* genes, a high-throughput, targeted sequence capture genotyping method was developed. Three PCR primer sets (Supporting Information Table S2) were designed to anneal to conserved regions of *KAI2d* genes and together were capable of amplifying *O. cernua KAI2d1-4* and *O. cumana KAI2d1-6* genes from parental lines and segregants using iProof High-Fidelity DNA Polymerase (BioRad #172530). The PCR products were then sequenced via MiSeq. Libraries were prepared using a Tn5 transposase which simultaneously fragments DNA to sizes less than 1000bp while ligating Nextera sequencing primers and indexing barcodes to each sample (Adey *et al*., 2010). DNA libraries were loaded onto MiSeq using an Illumina MiSeq Reagent Kit v3 (600 cycles) (Illumina, MS-102-3003) to generate 300bp paired-end reads. Over 30 million raw reads were generated by the MiSeq run. Data were split into fastq files based on indexing sequencing of each sample and Cutadaptor was used to trim Tn5 adaptor sequences (Martin, 2011). *KAI2d* genes were identified for each F_2_ plant using GATK to determine SNPs for each reference gene (McKenna *et al*., 2010). Raw reads were aligned to reference genes using BBMap software allowing up to one mismatch with zero gaps or substitutions (Bushnell, 2016). To determine whether a gene was present the alignments were checked manually for every sample using IGV Genome Browser (Robinson et al., 2011).

The genes *O. cernua KAI2d5-6* and *O. cumana KAI2d7-8* were not included in the high-throughput genotyping, so were genotyped by PCR followed by species-specific restriction enzyme digestion. DNA of each segregant was amplified and the resulting PCR product was digested and the products separated by agarose gel electrophoresis (see Supporting Information Table S3). PCRs used GoGreen Taq Plus Master Mix (Lamda Biotech) with general cycling conditions of 95 C for 2 min, 30 cycles of 90 C for 30 sec, 57 C for 30 sec (2 min for *OrceKAI2d6/ OrcuKAI2d8*), and 72 C for 30 sec, followed by 72 C for 5 min. Two assays were developed for distinguishing *OrceKAI2d5* and *OrcuKAI2d7* to provide a double check for PCR and restriction digestion efficiencies. Reliability of all assays was confirmed by including parental DNA as references in each run. In the event of ambiguous results or failure to amplify a product, the assay was repeated. Failure to amplify a product after three separate attempts in which positive controls were successful led to the conclusion that the gene was absent from the analyzed tissue. The DNA from all segregants was of adequate quality for PCR as judged by the fact that at least one of the four *KAI2d* genes was amplified from every segregant. One segregant was omitted due to insufficient DNA.

### Germination bio-assay for F_2_ phenotyping

F_2_ segregants were phenotyped based on percent germination of their F_3_ seeds in response to Oro, DCL or GR24 stimulants. Seeds were surface sterilized according to Plakhine *et al*. (2012) and 30-50 F_3_ seeds from a single F_2_ parent were sown on moist 9-mm glass fiber discs (Watman GFA), which were held in a Petri dish in darkness at 23°C for seven days. After conditioning, any spontaneous germination of seeds was recorded, the discs were blotted to remove water, and transferred to new Petri dishes. Each disc with seeds was considered an observation, with twelve disks distributed among three replicates of four treatments: 28 μL of water, DCL, Oro, or *rac-* GR24 at concentrations indicated above. The Petri dishes were then placed in 23°C for 10 days, after which seeds were scored for germination.

Two methods were used to correlate germination phenotypes with *KAI2* gene inheritance. First, a statistical model (described below) included genotype and phenotype data from all 94 F_2_ segregants. Second, a rationale was developed to exclude segregants with potentially distracting phenotypes as follows. Seeds of most segregants responded readily to GR24, but those that germinated in response to GR24 at a rate below 25% were also unresponsive to other stimulants, so were considered uninformative. Several other segregants showed higher than normal levels of spontaneous germination (details below), so were also disqualified. The remaining 77 segregants had a mean germination rate of 75% in response to GR24. These were classified into three main categories according to germination in response to a host-specific stimulant: 1) Oro, 2) DCL, and 3) Oro and DCL. Note that stimulants were applied singly in these assays. Segregants were further classified as high-response (greater than 40% germination rate) or low-response (39% or below), with a minimum of 8% over three replicates needed to be considered responsive to a stimulant. This minimum threshold was based on spontaneous germination in pure water, with most segregants showing no germination in such conditions. Several segregants with spontaneous germination rates above 10% responded to all stimulants and were removed from the genotyping analysis. Spontaneous germination in these lines has been reported previously (Plakhine *et al*., 2012), and likely involves signaling components aside from *KAI2* genes that contribute to low dormancy.

### Statistical modeling

A statistical model was developed to evaluate whether the germination responses of the 94 F_2_ segregants could be explained through the presence or absence of a single *KAI2d*, or a combination of *KAI2d* genes. We fit a Nested Generalized Linear Mixed-Effects Model (GLMM) using R (https://www.r-project.org/) and JAGS (Plummer, 2003) on the genotypes obtained through the targeted sequence capture assay. Through the GLMM, the number of seeds germinating under each experimental condition was modeled as a binomial random variate, with germination probability modeled through the Probit function as the additive effect of each *KAI2d* gene. Random effects were fit for Petri dishes nested within parent plants in order to account for Petri dish to Petri dish variation, as well as unmeasured characteristics of parent plants, genetic, and otherwise, termed ‘plant effects’.

A Bayesian approach was taken in parameter estimation (Hoff, 2009). In this case, we sought minimally informative prior distributions to conduct an objective analysis (Gelman, 2011). Marginal normal distributions were selected for fixed effects and marginal half normal for the random effects, both with mean 0 and a variance of 10. Priors for random effects variances are known to be more sensitive than priors for fixed effects (Fong *et al*., 2010), so we conducted a sensitivity analysis on the random effects prior by changing the prior variance hyper parameter to 1 and to 100 and found that it did not affect any of our conclusions.

Presence of *OrcuKAI2d3* and *OrcuKAI2d5* were found to be highly correlated 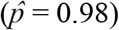, and therefore only their combined effect could be analyzed (the combined gene was marked as present if either *OrcuKAI2d3* or *OrcuKAI2d5* or both is present). JAGS Markov Chain Monte Carlo (MCMC) was found to converge by visual assessment of two chains: 400,000 sampling iterations were run after a 10,000 iteration burn in with a thinning rate of 100 to achieve 4,000 samples from the posterior for each quantity for each chain, and 8,000 in total posterior samples for each parameter. In order to assess significance, both practical and statistical, of model parameters, we produced symmetric 95% posterior credible intervals, as well as full posterior distributions approximated by MCMC.

### Functional complementation of *A. thaliana KAI2* mutants

RNA was isolated from seven-day conditioned seeds of *O. cernua* and *O. cumana* using the RNeasy method (Qiagen), with slight modifications (grinding in liquid nitrogen, 20% β-mercaptoethanol). *KAI2* coding sequences were amplified using RT-PCR on extracted RNA and cloned into pENTR/D-TOPO (ThermoFisher). Primer sequences are listed in Supporting Information Table S4. Entry clones were verified using Sanger sequencing and transferred into pKAI2pro-GW (a Gateway compatible vector containing the *A. thaliana KAI2* promoter) through Gateway recombination (Waters *et al*., 2015). Destination vectors were transformed into *Agrobacterium tumefaciens* strain GV3101, which was used to transform *A. thaliana* ecotype Landsberg *erecta kai2-2* mutants by the floral dip method (Clough & Bent, 1998). Transformed plants (T1), were selected on 0.5x Murashige-Skoog media supplemented with hygromycin (25μg/mL) (Harrison *et al*., 2006). The seed of T1 transgenic lines showing a segregation ratio of 3:1 hygromycin resistance were used in germination assays.

For each *KAI2* gene tested, 4-10 independent transformed *A. thaliana* lines were selected. The T1 plants were grown in 10-hours light/14-hours dark, at 22 °C for two weeks, then transferred to continuous light at 22 °C. Floral shoots were harvested under conditions favoring primary dormancy, when siliques were brown and the stem was green. The shoots were dried in paper bags at room temperature for three days, after which seeds were collected and stored at −80 °C until the germination assays.

Transgenic *A. thaliana* seeds were assayed for germination response. Seeds were surface sterilized for 2 minutes in 50% (v/v) bleach with 0.1% sodium dodecyl sulfate (w/v), rinsed with sterile deionized water 3 times, resuspended in 95% EtOH and immediately dried on sterile filter paper. These seeds were plated on 2.6 mM 2-(N-morpholino)ethanesulfonic acid (MES) media (pH 5.7), with 0.8% (w/v) BactoAgar, supplemented with either 0.1% acetone or 1 μM concentrations of the following stimulants: GR24, DCL, 5-deoxystrigol (5DS; StrigoLab) or Oro. Plates were placed under continuous light at 22 °C and germination was scored every 24 hrs for up to 5 days or until germination of the control exceeded 70%, whichever came first. Germination was defined as complete protrusion of the radical through the seed coat. Assays were replicated three times for each transgenic line and 4 - 10 independent seed lines were tested for each transgene.

Response to a stimulant was measured by mean germination percent. For each stimulant, mean germination percent was calculated by averaging the germination percent of 3 replicates, each including 30 seeds. Means, standard errors, and means comparisons using Tukey-Kramer HSD were performed using JMP software (JMP, Version 13.0, SAS Institute Inc.)

## Results

### Identification of putative stimulant receptor genes

We searched the *O. cernua* and *O. cumana* transcriptomes to identify transcripts of genes that are thought to be involved in SL perception. Single copies of *D14* and *MAX2* were identified within each transcriptome. The amino acid sequences of *D14* were found to be identical between the species, as they differed in just a single, silent nucleotide polymorphism. The *MAX2* sequences differed in two non-synonymous SNPs, which result in amino acid differences between the species (Supporting Information Fig. S1). However, comparison within a subset of F_2_ segregants differing in stimulant specificity indicated that neither amino acid change correlated with response to stimulant.

*KAI2* genes from each species were identified and characterized (Fig. 1). The *KAI2d* genes reported by Conn et al. (2015) as being evolved to respond to SLs were detected, comprising four genes in *O. cernua* and six in *O. cumana* (designated *OrceKAI2d1* to *d4* and *OrcuKAI2d1* to *d6*, respectively). Additionally, two previously unreported *KAI2* genes from each species were detected and named *OrceKAI2d5* and *d6* and *OrcuKAI2d7* and *d8* based on their grouping with the *KAI2d* genes rather than *KAI2i* or *KAI2c* genes (Fig. 1). One of these sequences, *OrceKAI2d6*, is likely a pseudogene as it is only about half the size of the full length *KAI2d* genes and includes stop codons. Thus, our analysis indicates the existence of 5 and 8 functional KAI2d proteins in *O. cernua* and *O. cumana*, respectively. Sequences of the conserved *KAI2* genes (*KAI2c* as defined by Conn et al., 2015) were also verified and included in the analysis of each species.

**Fig. 1.**
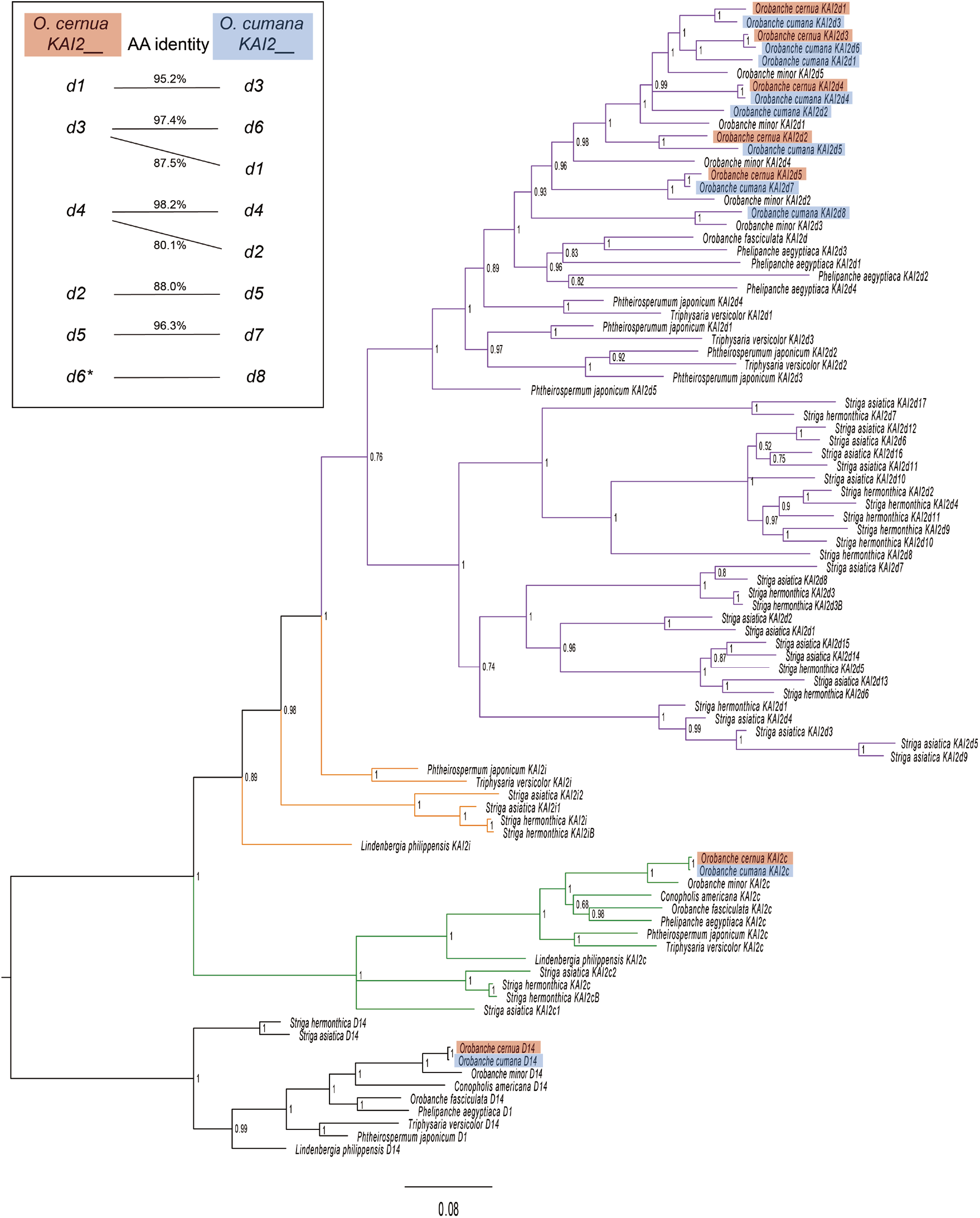
Relationship among *O. cernua* and *O. cumana KAI2* and *D14* genes (highlighted in red and blue, respectively) shown in the context of other known Orobanchaceae species. Bayesian phylogeny is based on coding sequences with *D14* as the outgroup. *KAI2* genes representing conserved (*KAI2c*), intermediate (*KAI2i*) and divergent (*KAI2d*) groups are indicated by different line colors. Posterior probabilities are indicated at nodes. Inset shows a schematic depiction of *O. cernua* and *O. cumana KAI2* putative paralog relationships. Percent amino acid (AA) identity is indicated between closest orthologues. \**O. cernua KAI2d6* is missing approximately half the coding region relative to the other genes, but aligns most closely with *O. cumana KAI2d8*. *OrceKAI2d6* is not included in the phylogenetic tree.

To validate the transcriptome assemblies, each *KAI2d* gene was cloned and sequenced from cDNA and genomic DNA to determine gene structures. All *KAI2d* genes share the same structure of two exons and one intron, with coding regions ranging in size from 810 to 825 bp and most introns ranging from 123 to 660 bp (the exception being *OrcuKAI2d8*, which exceeds 1 kb) (Supporting Information Table S5). Consistent with a recent common ancestry, *O. cernua* and *O. cumana* have several orthologous pairs of *KAI2d* genes, with amino acid identity ranging from 88.0 to 98.2%. However, gene duplication in the *O. cumana* lineage has produced additional paralogs: *OrcuKAI2d1* and *-d6* are most similar to *OrceKAI2d3*, while *OrcuKAI2d2* and -*d4* are most similar to *OrceKAI2d4* (Fig. 1 inset). All *KAI2* genes were expressed at all sampling times, although expression levels varied among the genes (Supporting Information Fig. S2). For *O. cernua*, the genes with highest expression were *OrceKAI2d4*, -*d5*, and -*d2*, whereas for *O. cumana*, the highest expression was for *OrcuKAI2d1*, -*d8*, and *-d4*. Given our hypothesis that key stimulant receptor genes are expressed only in a few cells near the micropyle (Plakhine *et al*., 2012), it is difficult to draw conclusions about the contribution of the observed gene expression levels to stimulant perception.

### Correlating *KAI2d* genotype with germination phenotype in segregants

*KAI2d* genotypes and germination response phenotypes were determined for 94 F_2_ individuals. F_2_ phenotypes were determined by assaying F_3_ seeds for responsiveness to Oro, DCL, and GR24 in individual assays. One approach to interpreting these results was to group segregants by their germination patterns after eliminating those that germinated at low rates or had high rates of spontaneous germination, leaving 77 F_2_ individuals with the clearest stimulant specificity. Most of these germinated in response to Oro, DCL, or both, but three segregants germinated in response to GR24, but not to the specific stimulants (Fig. 2). All F_3_ families that specifically responded to Oro contained the *OrceKAI2d2* gene, but also *OrceKAI2d1*, *OrceKAI2d4* and the pseudogene *OrceKAI2d6*, although these were not universally present. From the *O. cumana* side, many of the segregants contained *OrcuKAI2d2* and *OrcuKAI2d7*.

**Fig. 2.**
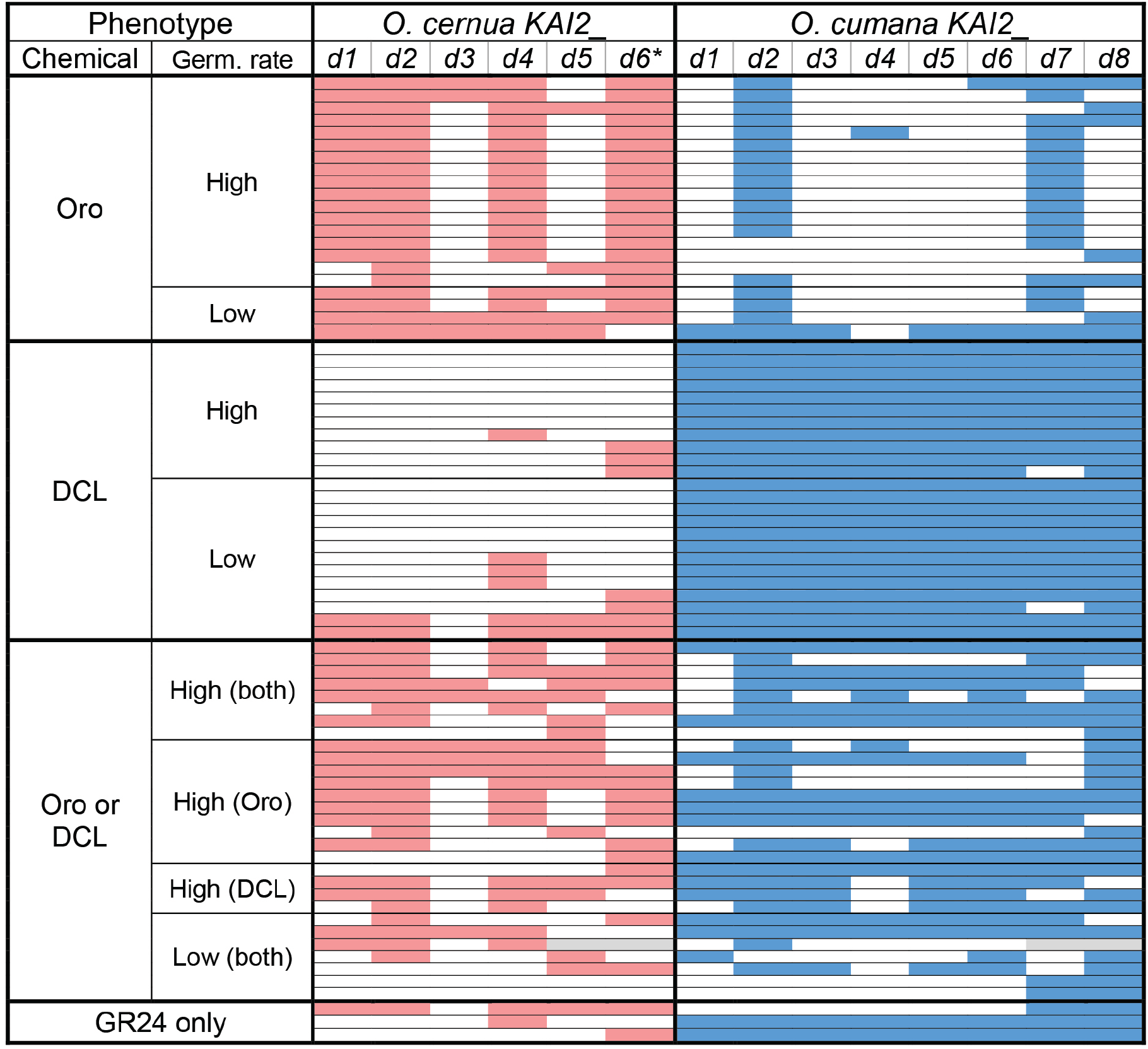
*KAI2d* gene presence for each *O. cernua* x *O. cumana* F_2_ segregants. Each of the 77 lines is indicated by a row and lines are grouped by phenotype category based on seed response to indicated stimulants. Filled cells indicate gene presence. Germination rates are defined as Low: 20-40%; High: >40%.

The pattern was reversed for F_3_ families that specifically responded to DCL, as these contained nearly all the *KAI2d* genes from *O. cumana*, but just a few from *O. cernua* (Fig. 2). Families that responded to both Oro and DCL showed diverse *KAI2d* gene compositions, generally containing multiple *KAI2d* genes from each parent, although a few families had just two genes detected. The most frequent *O. cernua* gene was again *OrceKAI2d2*, while *O. cumana* genes *OrcuKAI2d2* and *OrcuKAI2d8* were most represented from this parent. The final phenotype category consists of families that responded to GR24, but not to Oro or DCL. The pattern of *KAI2d* gene composition for these three families does not indicate a common theme.

We observed that the seeds of F_3_ families did not germinate at equal rates for different stimulants. For example, 74% of seeds from one line germinated in response to Oro while 34% germinated in response to DCL (data not shown). We thus divided germination phenotype into categories termed “low” (above background, but below 40%) and “high” (above 40%) to better evaluate whether specific genes or combinations of genes are associated with a qualitative change in germination rate. However, the patterns of *KAI2d* gene presence was similar for the high and low responders (Fig. 2).

Correlation analysis of *KAI2d* genotypes of the F_2_ individuals showed strong associations between *KAI2d* genes and suggest that they are linked in the *O. cernua* and *O. cumana* genomes (Fig. 3). This was evident in the apparent inheritance of *KAI2d* genes from each species in blocks (Fig. 2), which has likely complicated the genetic identification of any single gene as being responsible for conferring specificity to Oro or DCL.

**Fig. 3.**
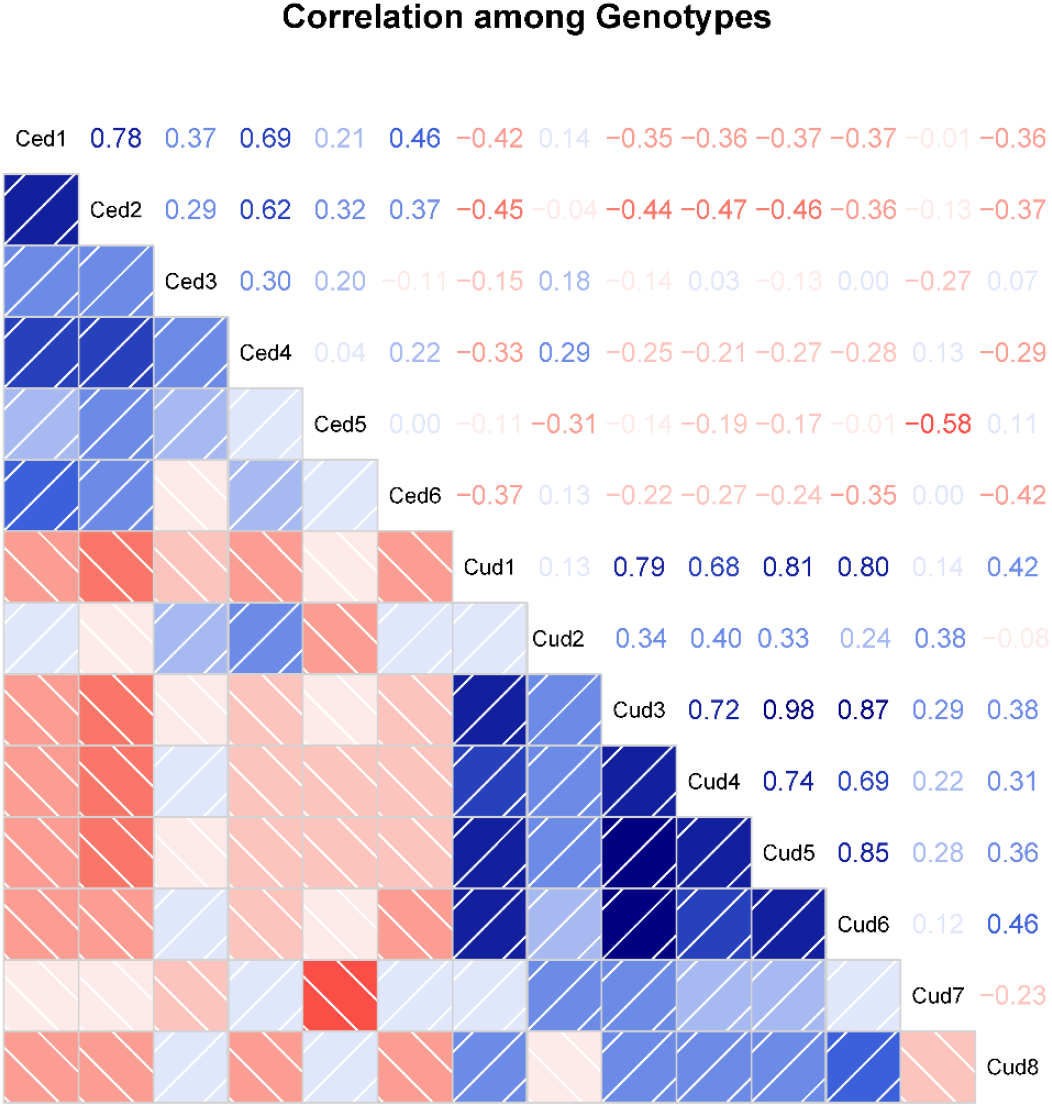
Corrgram showing the correlation among *KAI2d* genotypes. Darker blue indicates gene pairs are more highly correlated while darker red indicates less likelihood of correlation. Ce, *O. cernua;* Cu, *O. cumana*.

### Modeling germination rates and genotypes

To better understand the correlation between *KAI2d* genes and germination phenotype, we developed a statistical modeling strategy. For this we used the precise germination rates of all F_3_ families in response to each stimulant and statistically examined the effect of each *KAI2d* gene alone and in an additive manner, as well as the variance due to random effects. To assess significance of model parameters, symmetric 95% posterior credible intervals were calculated, as well as full posterior distributions approximated by MCMC for the effect of each gene on germination when exposed to each stimulant (Supporting Information Fig. S3, Table S6). Interpretation of model parameters in the Probit space follows the guideline that positive effects indicate that gene presence corresponds to higher germination rates, negative values the opposite, and magnitudes determine the strength of association. Thus, a gene is found to have a significant relationship if its posterior credible interval does not contain zero.

For the Oro response, the model strongly indicates that *OrceKAI2d2* co-occurs with increased germination, as indicated by the posterior distribution well above zero (Fig. 4, Supporting Information Table S6). To a lesser extent, *OrceKAI2d1* presence is associated with higher germination. None of the *O. cumana KAI2d* genes had a positive effect on germination, and *OrcuKAI2d1* and *OrcuKAI2d7* were associated with negative effects on Oro-induced germination. For the DCL response, genes with a significant credible interval for a positive effect on germination were *OrcuKAI2d3/5* (indistinguishable because of their tight linkage; see Fig. 3) and *OrcuKAI2d8*. One of the *O. cernua* genes, *OrceKAI2d1* was also associated with a positive effect on germination response to DCL. Genes that appear to have a negative effect on DCL-triggered germination include *OrcuKAI2d1* and *OrceKAI2d4* and *6*. No other genes were found to be correlated with increased or decreased germination rates (Supporting Information Fig. S3), but in all cases there was significant plant-to-plant as well as Petri dish-to-Petri dish variation. The “plant effect” suggests that there may be additional genetic influences on germination rates for DCL and Oro, such as other genes in the stimulant signaling pathway.

**Fig. 4.**
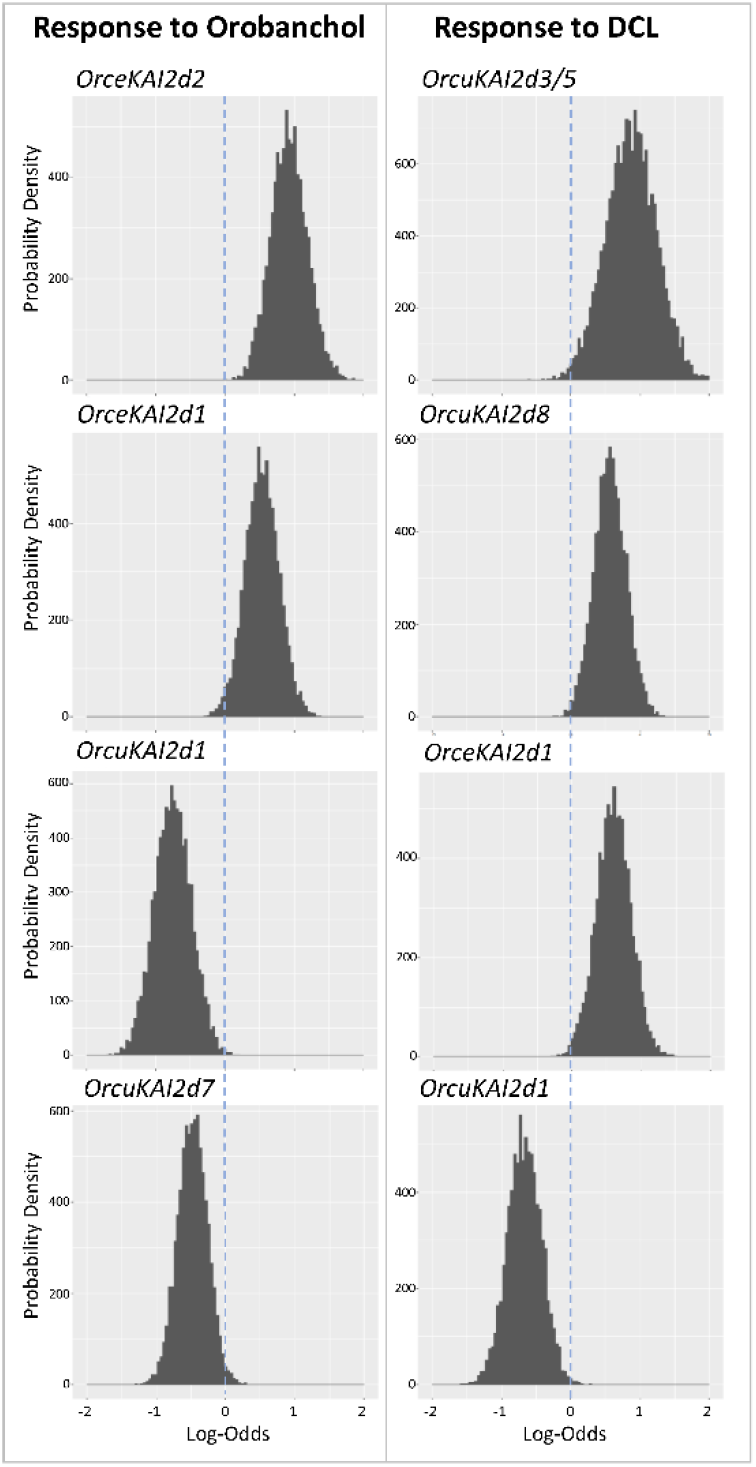
Posterior distribution charts for *KAI2d* genes in response to Orobanchol and DCL. The responses of selected specific genes to orobanchol (left column) and DCL (right column) are shown. Higher values of density indicate greater probability that the truth lies in that region, with peaks that do not include zero having the strongest correlation. Sign of log-odds indicates direction of correlation, and magnitude indicates strength of correlation. The dashed line indicates log-odds of zero in each chart. Additional charts are in Fig. S3.

### Assaying *KAI2* function in a cross-species complementation assay

Each of the 16 *O. cernua* and *O. cumana KAI2* genes (including KAI2c as well as all the KAI2d genes) were aligned to *AtKAI2* and *AtD14* and evaluated for the presence of amino acid residues known to be necessary for catalytic activity and interaction with *AtMAX2* (Supporting Information Fig. S4). Fifteen are predicted to encode functional proteins as they contain a full-length coding sequence, including the correct Ser95-His246-Asp217 catalytic triad needed for substrate hydrolysis (Zhao *et al*., 2015; Yao *et al*., 2016; Bythell-Douglas *et al*., 2017). The only exception is *OrceKAI2d6*, which is missing over two hundred and fifty bases and contains premature stop codons. The full-length proteins also conserve most of the amino acids identified as important in D14 protein association with MAX2/D3, notably the four residues considered essential (P161, E174, R177 and F180) as determined in *A. thaliana* (Yao *et al*., 2016).

*A. thaliana* does not germinate in the presence of natural SLs but is responsive to karrikins and the (-)-GR24 component of *rac*-GR24 (Scaffidi *et al*., 2014). The *A. thaliana kai2* mutant, which is insensitive to both treatments and shows increased seed dormancy compared to wild-type, has been used to evaluate specificities of parasitic plant *KAI2* genes through cross-species complementation assays (Conn *et al*., 2015; Toh *et al*., 2015). All *KAI2* cDNAs from *O. cernua* and *O. cumana* were cloned into a construct that drives expression from an *A. thaliana KAI2* promoter, and transformed into the *A. thaliana kai2-2* mutant background. The resulting transgenic *A. thaliana* plant lines were assayed for enhanced germination in response to SLs (Oro, 5DS and *rac*-GR24) and DCL. The non-functional *OrceKAI2d6* and an empty vector control were included as negative controls (Supporting Information Fig. S5).

Only one *O. cernua* gene, *OrceKAI2d2*, was able to consistently confer enhanced germination in response to SLs in the *kai2-2* mutant background (Fig. 5, Table 1). The homolog of this gene in *O. cumana, OrcuKAI2d5*, produced the same phenotype, suggesting similar specificities in response to SLs. Interestingly, the *O. cumana* genes *OrcuKAI2d7* and *OrcuKAI2d8* also complemented the mutant in response to SLs, whereas the corresponding *O. cernua* homologs did not. None of the *KAI2* genes tested were found to consistently and specifically confer a germination response to DCL (Table 1, Supporting Information Fig. S5). In one instance the *OrcuKAI2d3* gene conferred DCL responsiveness to an *A. thaliana* line, but the effect was non-specific and was not repeated in nine other independently transformed lines.

**Fig. 5.**
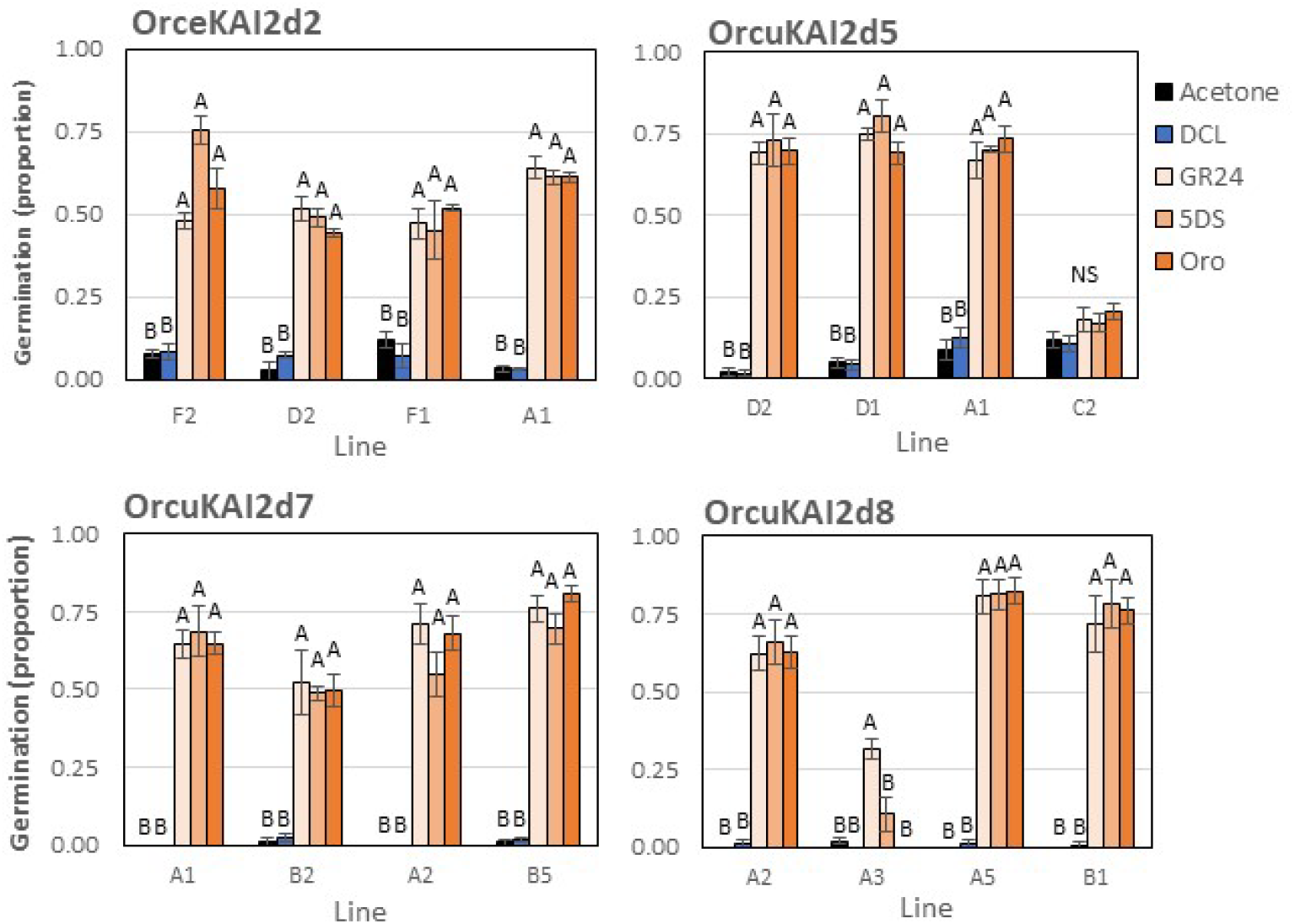
Assays of *KAI2d* gene responses to germination stimulants in a heterologous system. The indicated genes were expressed as transgenes in the *A. thaliana kai2* mutant background. Seeds were scored for germination following exposure to acetone (negative control), DCL, GR24, 5DS or Oro. Each line represents a unique transformation event. Tukey-Kramer HSD test was used to determine significance, * P < 0.05. Each column represents the mean of 3 replications and vertical bracket represent SE.

**Table 1.**
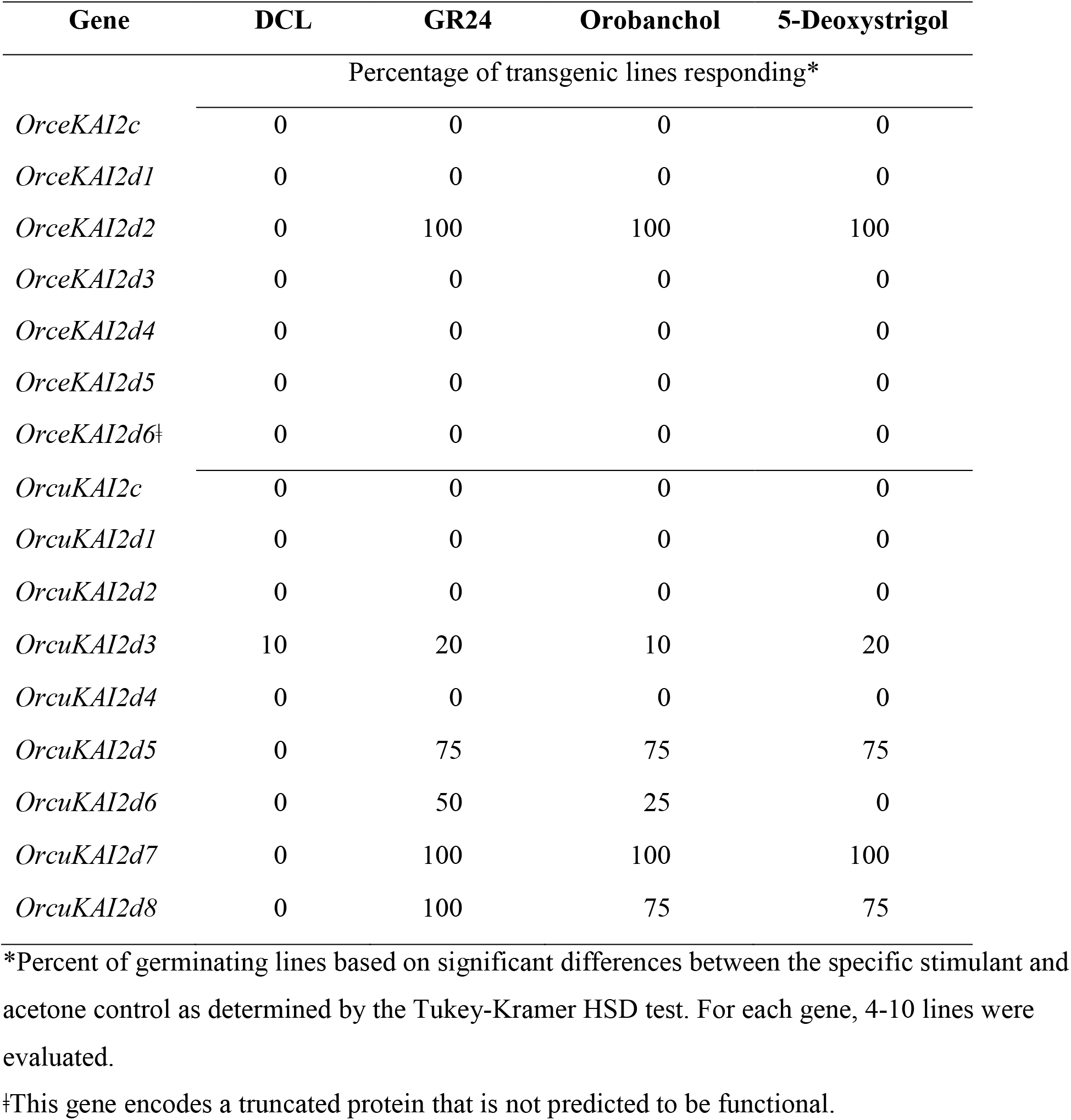
Summary of response of each *Orobanche KAI2* transgene expressed in the *Arabidopsis kai2* mutant background and assayed by determining germination in response to DCL and SL germination stimulants.

## Discussion

All data indicate that *OrceKAI2d2* is a SL receptor in *O. cernua*. This gene segregated with Oro responsiveness in the analysis of *O. cernua* x *O. cumana* F_3_ families (Figs. 2, 4), and was able to complement the *kai2-2* mutant phenotype when treated with Oro and 5DS in the heterologous assay system (Fig. 5). Finding this association in *O. cernua*, which commonly attacks Oro-producing tomato hosts is not surprising, although is it interesting that just one of the five putatively functional *O. cernua KAI2d* genes was clearly indicated. This somewhat parallels findings from *Striga*, which contains multiple *KAI2/HTL* genes, with 13 reported in *S. hermonthica* (Conn *et al*., 2015) and 17 in *S. asiatica* (Yoshida *et al*., 2019). Of the *S. hermonthica* genes, at least six were found to respond to SLs in a heterologous assay, but just one (*ShHTL7*) was extraordinarily sensitive to SLs (Toh *et al*., 2015). The apparent linkage of *KAI2d* genes on *Orobanche* chromosomes (Fig. 3) complicates the ability to distinguish the effects of other *KAI2d* genes, but is consistent with findings of linked *KAI2d* genes in *S. asiatica* (Yoshida *et al*., 2019). Our statistical model suggests a weaker contribution of *OrceKAI2d1* to SL germination response, although this was not supported by the heterologous assay (Table 1). We note that a lack of response to SLs in the heterologous assay system does not necessarily preclude a *KAI2d* gene as a functional SL receptor in its native context. In some cases, KAI2d proteins from *Orobanche* spp. may not be stably expressed or able to interact sufficiently well with *Arabidopsis* MAX2 and SMAX1 signaling partners to promote germination.

The *O. cumana* germination response to DCL from sunflower is not clearly explained by the current data. The strong association of strict DCL response with inheritance of (in most cases) all eight *O. cumana KAI2d* genes argues for involvement of a gene or regulatory element associated with that block of genes (Fig. 2). Despite *O. cumana* having an expanded set of *KAI2d* genes (Fig. 1) that would seem to be ideal for evolving new substrate specificities, none of the *O. cumana* genes responded to DCL in the heterologous assay (Table 1). Rather, this approach indicates that *O. cumana* has retained genes that are capable of SL-response. *OrcuKAI2d5*, the ortholog of SL-responsive *OrceKAI2d2*, was able to complement the *kai2* mutant phenotype in response to Oro and 5DS, indicating that it has conserved the ability to bind and transduce a SL signal. Additionally, the *O. cumana* genes *OrcuKAI2d7* and *OrcuKAI2d8* responded to SLs in the heterologous assay (Fig. 5, Table 1). It is interesting that the *O. cumana* genes produced a response, whereas the *O. cernua* homologs *OrceKAI2d5* and *OrceKAI2d6* showed no complementation, although the lack of SL response by *OrceKAI2d6* was expected because it encodes a truncated protein. The lack of response to SLs by *OrceKAI2d5* may suggest a divergence from its *O. cumana* homolog or a limitation of the assay system. But taken together, the finding that *O. cumana* contains *KAI2d* genes that respond to Oro or 5DS, while seeds of *O. cumana* do not germinate in response to these molecules, suggests that other factors are involved in regulating germination.

This work fits the current model of SL germination signaling perception in parasitic plants with respect to *O. cernua*, but raises additional questions. One issue is the failure to explain why *O. cumana* seeds do not germinate in response to SLs despite having at least three forms of *KAI2d* that respond to Oro in a heterologous assay. It is possible that the SL perception pathway in *O. cumana* has been modified with respect to germination. One postulated mechanism is that KAI2d proteins could be flipped to function as germination inhibitors if their SL biding capacities were retained while mutations occurred that reduce interactions with binding partners such as MAX2 or SMAX1 (Nelson, 2021). Such stabilization of an incomplete multiprotein assembly could competitively enhance retention of SMAX1 and maintain inhibition of germination. Other possible mechanisms include limiting expression of SL-responsive *KAI2* genes during conditioning (notably, *OrcuKAI2d5* is among the genes with lowest expression in the transcriptomes; Fig. S2) or spatially restricting *KAI2d* expression to key cells in the parasite seed that respond to stimulants (Plakhine *et al*., 2012; Tsuchiya *et al*., 2015).

The other persistent question is how *O. cumana* can germinate in response to DCL when none of its *KAI2d* genes directly respond to DCL in the heterologous assay. Failure to identify the DCL receptor could be explained by the presence of an additional DCL receptor that is not part of the *KAI2d* family, although our correlation analysis indicates that the putative DCL-responsive element segregates with the *KAI2d* genes. It is also possible that the complementation assay in *A. thaliana* has limitations and does not precisely mimic germination signaling in *Orobanche* seeds. Despite excellent studies of parasite KAI2d substrate specificities (Yao *et al*., 2017; de Saint Germain *et al*., 2021), such interactions may not translate completely to predict responses by parasite seeds. For example, it is conceivable that DCL undergoes metabolism in *O. cumana*, but not *A. thaliana*, to produce a ligand that can be perceived by one of the *OrcuKAI2d* proteins. Unlike SLs, DCL does not have a cleavable methylbutenolide D-ring, which has been associated with activation of SL receptors (de Saint Germain *et al*., 2016; Yao *et al*., 2017; Uraguchi *et al*., 2018). In this way DCL resembles Karrikins, which also lack a cleavable D-ring yet they – or an undefined metabolite – signal germination through KAI2 in many nonparasitic plant species (Yao & Waters, 2020).

In summary, this study supports a model in which *KAI2d* proteins recognize SL molecules as part of the regulation of germination in parasitic plant seeds. A combination of genetic correlations and heterologous complementation studies strongly point to *OrceKAI2d2* as a primary receptor for the SL stimulant Oro in *O. cernua*. A strong candidate receptor for perception of DCL was not identified, but the chromosomal region(s) containing the KAI2d genes should be targeted for future research. These data further suggest that regulation of germination specificity is more complicated than the simple presence or absence of any individual *KAI2d* gene (as suggested by others, e.g., Brun *et al*., 2020). *KAI2d* proteins with affinity for SLs must function in coordination with other factors that contribute to germination in the context of a parasite seed. More research is needed on *KAI2* genes – and the entire signaling pathway – to understand regulation of germination response to SLs and other classes of environmental signaling molecules.

## Acknowledgements

This work was funded by BARD award no. US-4616-13 to JHW, HE, and YT, with additional support from BARD Graduate Student Travel Award GS-33-2016 (HL), the US National Institute of Food and Agriculture 135997 (JHW), and NSF IOS-1856741 (DCN). We thank Dr. Koichi Yoneyama for providing orobanchol and Robert Tuosto for technical assistance, Dr. Jacob Kitzman (University of Michigan) for use of a MiSeq, and Dr. David Haak for advice.

## Author contributions

HE, DMJ, YT and JHW conceived and supervised the project. DP generated the segregating population. HE and YT isolated RNA from parental lines, and HL isolated genomic DNA from parent and their segregants. HL and NZ genotyped the segregants. NW developed the statistical model in coordination with HL. CC performed the phylogenetic analysis. CC, DCN, and HL cloned KAI2 genes. DCN advised on complementation assays and data interpretation. HL conducted bioinformatics analyses, designed the KAI2 sequencing strategy, and performed the complementation assay. HL wrote the manuscript. JHW edited the manuscript with input from all co-authors.

## Data availability

All data are available at NCBI. Transcriptome raw reads and assemblies are in the GEO repository number GSE196954. Specific *KAI2* gene sequences are at NCBI, with accessions OMB837823 - OMB837829 for *OrceKAI2c* and *OrceKAI2d1-6* and OMB811642 - OMB811650 for *OrcuKAI2c* and *OrcuKAI2d1-8*.

## References

Adey A, Morrison HG, Asan, Xun X, Kitzman JO, Turner EH, Stackhouse B, MacKenzie AP, Caruccio NC, Zhang X, Shendure J. 2010. Rapid, low-input, low-bias construction of shotgun fragment libraries by high-density in vitro transposition. Genome Biology 11: R119.

Altschul SF, Madden TL, Schäffer AA, Zhang J, Zhang Z, Miller W, Lipman DJ. 1997. Gapped BLAST and PSI-BLAST: a new generation of protein database search programs. Nucleic Acids Research 25: 3389–3402.

Andrews S. 2010. FastQC a quality control tool for high throughput sequence data. http://www.bioinformatics.babraham.ac.uk/projects/fastqc/. [accessed 2018].

Bolger AM, Lohse M, Usadel B. 2014. Trimmomatic: a flexible trimmer for Illumina sequence data. Bioinformatics 30: 2114–2120.

Brun G, Spallek T, Simier P, Delavault P. 2020. Molecular actors of seed germination and haustoriogenesis in parasitic weeds. Plant Physiology 185: 1270–1281.

Bunsick M, Toh S, Wong C, Xu Z, Ly G, McErlean CSP, Pescetto G, Nemrish KE, Sung P, Li JD, Scholes JD, Lumba S. 2020. SMAX1-dependent seed germination bypasses GA signalling in *Arabidopsis* and *Striga*. Nature Plants 6: 646–652.

Burger M, Chory J. 2020. The many models of strigolactone signaling. Trends in Plant Science 25: 395–405.

Bushnell B. 2016. BBMap short read aligner. https://sourceforge.net/projects/bbmap/. [accessed 2018].

Bythell-Douglas R, Rothfels CJ, Stevenson DWD, Graham SW, Wong GK-S, Nelson DC, Bennett T. 2017. Evolution of strigolactone receptors by gradual neo-functionalization of KAI2 paralogues. BMC Biol. 15: 52.

Clough SJ, Bent AF. 1998. Floral dip: a simplified method for *Agrobacterium-mediated* transformation of *Arabidopsis thaliana*. Plant Journal 16: 735–743.

Conn CE, Bythell-Douglas R, Neumann D, Yoshida S, Whittington B, Westwood JH, Shirasu K, Bond CS, Dyer KA, Nelson DC. 2015. Convergent evolution of strigolactone perception enabled host detection in parasitic plants. Science 349: 540–543.

Conn CE, Nelson DC. 2016. Evidence that KARRIKIN-INSENSITIVE2 (KAI2) receptors may perceive an unknown signal that is not karrikin or strigolactone. Frontiers in Plant Science 6.

De Cuyper C, Struk S, Braem L, Gevaert K, De Jaeger G, Goormachtig S. 2017. Strigolactones, karrikins and beyond. Plant Cell Environ. 40: 1691–1703.

de Saint Germain A, Clavé G, Badet-Denisot M-A, Pillot J-P, Cornu D, Le Caer J-P, Burger M, Pelissier F, Retailleau P, Turnbull C, Bonhomme S, Chory J, Rameau C, Boyer F-D. 2016. An histidine covalent receptor and butenolide complex mediates strigolactone perception. Nature Chemical Biology 12: 787–794.

de Saint Germain A, Jacobs A, Brun G, Pouvreau J-B, Braem L, Cornu D, Clavé G, Baudu E, Steinmetz V, Servajean V, Wicke S, Gevaert K, Simier P, Goormachtig S, Delavault P, Boyer F-D. 2021. A *Phelipanche ramosa* KAI2 protein perceives strigolactones and isothiocyanates enzymatically. Plant Communications: 100166.

Dor E, Yoneyama K, Wininger S, Kapulnik Y, Yoneyama K, Koltai H, Xie XN, Hershenhorn J. 2011. Strigolactone deficiency confers resistance in tomato line SL-ORT1 to the parasitic weeds *Phelipanche* and *Orobanche* spp. Phytopathology 101: 213–222.

Doyle JJ, Doyle JL. 1987. A rapid DNA isolation procedure for small quantities of fresh leaf tissue.

Fong Y, Rue H, Wakefield J. 2010. Bayesian inference for generalized linear mixed models. Biostatistics 11: 397–412.

Gelman A. 2011. Induction and deduction in Bayesian data analysis. Rationality, Markets and Morals 2: 67–78.

Haas BJ, Papanicolaou A, Yassour M, Grabherr M, Blood PD, Bowden J, Couger MB, Eccles D, Li B, Lieber M, MacManes MD, Ott M, Orvis J, Pochet N, Strozzi F, Weeks N, Westerman R, William T, Dewey CN, Henschel R, LeDuc RD, Friedman N, Regev A. 2013. *De novo* transcript sequence reconstruction from RNA-seq using the Trinity platform for reference generation and analysis. Nature Protocols 8: 1494–1512.

Harrison SJ, Mott EK, Parsley K, Aspinall S, Gray JC, Cottage A. 2006. A rapid and robust method of identifying transformed Arabidopsis thaliana seedlings following floral dip transformation. Plant Methods 2: 19.

Hegenauer V, Körner M, Albert M. 2017. Plants under stress by parasitic plants. Current Opinion in Plant Biology 38: 34–41.

Hoff P. 2009. A first course in Bayesian statistical methods.. New York: SpringerVerlag.

Ishida JK, Wakatake T, Yoshida S, Takebayashi Y, Kasahara H, Wafula E, dePamphilis CW, Namba S, Shirasu K. 2016. Local auxin biosynthesis mediated by a YUCCA flavin monooxygenase regulates haustorium development in the parasitic plant *Phtheirospermum japonicum*. Plant Cell 28: 1795–1814.

Joel DM, Bar H, Mayer AM, Plakhine D, Ziadne H, Westwood JH, Welbaum GE. 2011a. Seed ultrastructure and water absorption pathway of the root-parasitic plant *Phelipanche aegyptiaca* (Orobanchaceae). Annals of Botany 109: 181–195.

Joel DM, Chaudhuri SK, Plakhine D, Ziadna H, Steffens JC. 2011b. Dehydrocostus lactone is exuded from sunflower roots and stimulates germination of the root parasite *Orobanche cumana*. Phytochemistry 72: 624–634.

Khosla A, Morffy N, Li Q, Faure L, Chang SH, Yao J, Zheng J, Cai ML, Stanga J, Flematti GR, Waters MT, Nelson DC. 2020. Structure–function analysis of SMAX1 reveals domains that mediate its karrikin-induced proteolysis and interaction with the receptor KAI2. The Plant cell 32: 2639–2659.

Koltai H. 2014. Receptors, repressors, PINs: a playground for strigolactone signaling. Trends in Plant Science 19: 727–733.

Langmead B, Salzberg SL. 2012. Fast gapped-read alignment with Bowtie 2. Nature Methods 9: 357–U354.

Machin DC, Hamon-Josse M, Bennett T. 2020. Fellowship of the rings: a saga of strigolactones and other small signals. New Phytologist 225: 621–636.

Martin M. 2011. Cutadapt removes adapter sequences from high-throughput sequencing reads. EMBnet J 17: 10 – 12.

McKenna A, Hanna M, Banks E, Sivachenko A, Cibulskis K, Kernytsky A, Garimella K, Altshuler D, Gabriel S, Daly M, DePristo MA. 2010. The Genome Analysis Toolkit: a MapReduce framework for analyzing next-generation DNA sequencing data. Genome research 20: 1297–1303.

Nelson DC. 2021. The mechanism of host-induced germination in root parasitic plants. Plant Physiology 185: 1353–1373.

Parker C. 2009. Observations on the current status of *Orobanche* and *Striga* problems worldwide. Pest Man. Sci. 65: 453–459.

Parker C. 2012. Parasitic weeds: A world challenge. Weed Sci. 60: 269–276.

Parker C. 2013. The parasitic weeds of the Orobanchaceae. In: Joel DM, Gressel J, Musselman LJ eds. Parasitic Orobanchaceae: Springer Berlin Heidelberg, 313–344.

Parker C, Riches CR. 1993. Parasitic Weeds of the World: Biology and Control. Wallingford: CAB International.

Parra G, Bradnam K, Korf I. 2007. CEGMA: a pipeline to accurately annotate core genes in eukaryotic genomes. Bioinformatics 23: 1061–1067.

Plakhine D, Tadmor Y, Ziadne H, Joel DM. 2012. Maternal tissue is involved in stimulant reception by seeds of the parasitic plant *Orobanche*. Annals of Botany 109: 979–986.

Plummer M. 2003. JAGS: A program for analysis of Bayesian graphical models using Gibbs sampling. Proceedings of the 3rd International Workshop on Distributed Statistical Computing (DSC 2003). 20–22.

Ronquist F, Teslenko M, van der Mark P, Ayres DL, Darling A, Höhna S, Larget B, Liu L, Suchard MA, Huelsenbeck JP. 2012. MrBayes 3.2: Efficient Bayesian phylogenetic Inference and model choice across a large model space. Systematic Biology 61: 539–542.

Samejima H, Babiker AG, Takikawa H, Sasaki M, Sugimoto Y. 2016. Practicality of the suicidal germination approach for controlling *Striga hermonthica*. Pest Management Science 72: 2035–2042.

Scaffidi A, Waters MT, Sun YK, Skelton BW, Dixon KW, Ghisalberti EL, Flematti GR, Smith SM. 2014. Strigolactone hormones and their stereoisomers signal through two related receptor proteins to induce different physiological responses in Arabidopsis. Plant Physiology 165: 1221–1232.

Spallek T, Melnyk CW, Wakatake T, Zhang J, Sakamoto Y, Kiba T, Yoshida S, Matsunaga S, Sakakibara H, Shirasu K. 2017. Interspecies hormonal control of host root morphology by parasitic plants. Proc. Natl. Acad. Sci. USA.

Stanga JP, Smith SM, Briggs WR, Nelson DC. 2013. SUPPRESSOR OF MORE AXILLARY GROWTH2 1 controls seed germination and seedling development in Arabidopsis. Plant Physiology 163: 318–330.

Toh S, Holbrook-Smith D, Stogios PJ, Onopriyenko O, Lumba S, Tsuchiya Y, Savchenko A, McCourt P. 2015. Structure-function analysis identifies highly sensitive strigolactone receptors in *Striga*. Science 350: 203–207.

Tsuchiya Y, Yoshimura M, Sato Y, Kuwata K, Toh S, Holbrook-Smith D, Zhang H, McCourt P, Itami K, Kinoshita T, Hagihara S. 2015. Probing strigolactone receptors in *Striga hermonthica* with fluorescence. Science 349: 864–868.

Umehara M, Hanada A, Yoshida S, Akiyama K, Arite T, Takeda-Kamiya N, Magome H, Kamiya Y, Shirasu K, Yoneyama K, Kyozuka J, Yamaguchi S. 2008. Inhibition of shoot branching by new terpenoid plant hormones. Nature 455: 195–200.

Uraguchi D, Kuwata K, Hijikata Y, Yamaguchi R, Imaizumi H, AM S, Rakers C, Mori N, Akiyama K, Irle S, McCourt P, Kinoshita T, Ooi T, Tsuchiya Y. 2018. A femtomolar-range suicide germination stimulant for the parasitic plant *Striga hermonthica*. Science 362: 1301–1305.

Wang L, Xu Q, Yu H, Ma HY, Li XQ, Yang J, Chu JF, Xie Q, Wang YH, Smith SM, Li JY, Xiong GS, Wang B. 2020. Strigolactone and karrikin signaling pathways elicit ubiquitination and proteolysis of SMXL2 to regulate hypocotyl elongation in Arabidopsis. Plant Cell 32: 2251–2270.

Waters MT. 2017. From little things big things grow: karrikins and new directions in plant development. Functional Plant Biology 44: 373–385.

Waters MT, Scaffidi A, Moulin SLY, Sun YK, Flematti GR, Smith SM. 2015. A *Selaginella moellendorffii* ortholog of KARRIKIN INSENSITIVE2 functions in Arabidopsis development but cannot mediate responses to karrikins or strigolactones. The Plant cell 27: 1925.

Waters MT, Smith SM. 2013. KAI2- and MAX2-mediated responses to karrikins and strigolactones are largely independent of HY5 in Arabidopsis seedlings. Molecular Plant 6: 63–75.

Westwood JH, Yoder JI, Timko MP, dePamphilis CW. 2010. The evolution of parasitism in plants. Trends Plant Sci. 15: 227–235.

Wigchert SCM, Kuiper E, Boelhouwer GJ, Nefkens GHL, Verkleij JAC, Zwanenburg B. 1999. Dose-response of seeds of the parasitic weeds *Striga* and *Orobanche* toward the synthetic germination stimulants GR 24 and Nijmegen 1. Journal of Agricultural and Food Chemistry 47: 1705–1710.

Yang T, Lian YK, Wang CY. 2019. Comparing and contrasting the multiple roles of butenolide plant growth regulators: strigolactones and karrikins in plant development and adaptation to abiotic stresses. International journal of molecular sciences 20: 36.

Yao JR, Waters MT. 2020. Perception of karrikins by plants: a continuing enigma. Journal of Experimental Botany 71: 1774–1781.

Yao R, Ming Z, Yan L, Li S, Wang F, Ma S, Yu C, Yang M, Chen L, Chen L, Li Y, Yan C, Miao D, Sun Z, Yan J, Sun Y, Wang L, Chu J, Fan S, He W, Deng H, Nan F, Li J, Rao Z, Lou Z, Xie D. 2016. DWARF14 is a non-canonical hormone receptor for strigolactone. Nature 536: 469–473.

Yao R, Wang F, Ming Z, Du X, Chen L, Wang Y, Zhang W, Deng H, Xie D. 2017. ShHTL7 is a non-canonical receptor for strigolactones in root parasitic weeds. Cell Research 27: 838.

Yoneyama K, Awad AA, Xie X, Yoneyama K, Takeuchi Y. 2010. Strigolactones as germination stimulants for root parasitic plants. Plant and Cell Physiology 51: 1095–1103.

Yoneyama K, Ruyter-Spira C, Bouwmeester H. 2013. Induction of germination. In: Joel DM, Gressel J, Musselman LJ eds. Parasitic Orobanchaceae: Springer Berlin Heidelberg, 167–194.

Yoshida S, Kim S, Wafula EK, Tanskanen J, Kim Y-M, Honaas L, Yang Z, Spallek T, Conn CE, Ichihashi Y, Cheong K, Cui S, Der JP, Gundlach H, Jiao Y, Hori C, Ishida JK, Kasahara H, Kiba T, Kim M-S, Koo N, Laohavisit A, Lee Y-H, Lumba S, McCourt P, Mortimer JC, Mutuku JM, Nomura T, Sasaki-Sekimoto Y, Seto Y, Wang Y, Wakatake T, Sakakibara H, Demura T, Yamaguchi S, Yoneyama K, Manabe R-i, Nelson DC, Schulman AH, Timko MP, dePamphilis CW, Choi D, Shirasu K. 2019. Genome sequence of *Striga asiatica* provides insight into the evolution of plant parasitism. Current Biology 29: 3041–3052.e3044.

Zhao L-H, Zhou XE, Yi W, Wu Z, Liu Y, Kang Y, Hou L, de Waal PW, Li S, Jiang Y, Scaffidi A, Flematti GR, Smith SM, Lam VQ, Griffin PR, Wang Y, Li J, Melcher K, Xu HE. 2015. Destabilization of strigolactone receptor DWARF14 by binding of ligand and E3-ligase signaling effector DWARF3. Cell Research 25

