## Supplemental tables and figures for "An *Orobanche cernua* x *Orobanche cumana* segregating population provides insight into the regulation of germination specificity in a parasitic plant"

**Table S1**. *Orobanche cernua* and *O. cumana* transcriptome statistics.

| | *O. cernua* | *O. cumana* |
|---|---|---|
| Raw Reads | 391,787,707 | 433,836,952 |
| Number of assembled contigs | 204,992 | 216,881 |
| N50 | 1,631 | 1,572 |
| Mean contig length | 940.75 | 895.6 |
| Total assembled bases | 192,845,756 | 194,239,216 |
| Mapping % of input reads (Bowtie2) | 97.56% | 97.70% |
| Predicted ESTs | 103,570 | 110,019 |

**Table S2.** Universal primer sequences used to amplify *OrceKAI2d1-4* and *OrcuKAI2d1-6* for the targeted sequence capture assay.

| Primer ID | Sequence | Genes Amplified - *O. cernua* | Genes Amplified - *O. cumana* |
|---|---|---|---|
| R1 | TCATGCATCAACAATATC | | *OrcuKAI2d5, OrcuKAI2d3, OrcuKAI2d2* |
| R2 | TCAACCATCAACAATATC | *OrceKAI2d4* | *OrcuKAI2d4* |
| R3 | TCAGGCAGCGATATTATA | *OrceKAI2c* | *OrcuKAI2c* |
| R4 | TCATGCATCAATATCGTG | *OrceKAI2d3* | *OrcuKAI2d6* |
| F1 | ATGGGAATCACCCAAG | *OrceKAI2c* | *OrcuKAI2c* |
| F2 | ATGAACCGTATAGTTGGACT | | *OrcuKAI2d5* |
| F3 | ATGAGTAGCATAGTTGGTG | *OrceKAI2d1, OrceKAI2d4, OrceKAI2d3* | *OrcuKAI2d1, OrcuKAI2d3, OrcuKAI2d4, OrcuKAI2d6* |
| F2B | ATGAACAGCATAGTTGGACT | *OrceKAI2d2* | |
| F3B | ATGGGTAGCATTGTTG | | *OrcuKAI2d2* |
| R1B | TCATACATCAGCAATATC | *OrceKAI2d2* | |

**Table S3**. Genotyping assays for genes *OrceKAI2d5* & *6* and *OrcuKAI2d7* & *8*. DNA from each segregant was amplified by PCR using primers specific to a pair of genes, and then the products were digested with restriction enzymes specific to the species. Conclusions about the presence of a given gene was made based on the number and size of the bands visualized by gel electrophoresis.

| Gene names | Primers | Restriction enzyme | Band sizes | Conclusion |
|---|---|---|---|---|
| *OrceKAI2d5* and *OrcuKAI2d7* (Assay 1) | F: ATGAACATAGTTGGAGCAGC<br>R: CGTTTCCATCGCGTCTAGGA | SalI | 600bp | *OrceKAI2d5* |
| | | | 120bp, 480bp | *OrcuKAI2d7* |
| *OrceKAI2d5* and *OrcuKAI2d7* (Assay 2) | F: ACTCAACCCTAGAAGGCCAC<br>R: CGTTTCCATCGCGTCTAGGA | MseI | 100bp, 120bp, 170bp | *OrceKAI2d5* |
| | | | 170bp, 220bp | *OrcuKAI2d7* |
| *OrceKAI2d6* and *OrcuKAI2d8* | F: ATGAGCACAGTTGGAGC<br>R: TCAGGTCGGTATATCGAAGCC | SphI | 320bp | *OrceKAI2d6* |
| | | | 100bp, 220bp | *OrcuKAI2d8* |

**Table S4**. Primer sequences used for amplifying *KAI2d* sequences for pENTR/D-TOPO cloning.

| gene | F primer | R primer |
|---|---|---|
| *OrceKAI2c* | CACCATGGGAATCACCCAAGACGCT | TCAGGCAGCGATATTATAAC |
| *OrceKAI2d1* | CACCATGAGTAGCATAGTTGGTGCC | TCATGCATCAACAATACCGA |
| *OrceKAI2d2* | CACCATGAACAGCATAGTTGGACTT | TCATACATCAGCAATATCGC |
| *OrceKAI2d3* | CACCATGAGTAGCATAGTTGGTGCC | TCATGCATCAATATCGTGAT |
| *OrceKAI2d4* | CACCATGAGTAGCATAGTTGGTGCG | TCAACCATCAACAATATCGT |
| *OrceKAI2d5* | CACCATGAACATAGTTGGAGCA | TCAGGCGTCAATGATGTC |
| *OrceKAI2d6* | CACCATGAGCACAGTTGGAGC | TCAGGCTATATCGTGTTGTAT |
| *OrcuKAI2c* | CACCATGGGAATCACCCAAGAAGCT | TCAGGCAGCGATATTATAAC |
| *OrcuKAI2d1* | CACCATGAGTAGCATAGTTGGTGCC | ATCGTGCCCCCGGCATACT |
| *OrcuKAI2d2* | CACCATGGGTAGCATTGTTGGTGCG | TCATGCATCAACAATATCAT |
| *OrcuKAI2d3* | CACCATGAGTAGCATAGTTGGTGCC | TCATGCATCAACAATATCGA |
| *OrcuKAI2d4* | CACCATGAGTAGCATAGTTGGTGCG | TCAACCATCAACAATATCGT |
| *OrcuKAI2d5* | CACCATGAACCGTATAGTTGGACTT | TCATGCATCAACAATATCGC |
| *OrcuKAI2d6* | CACCATGAGTAGCATAGTTGGTGCC | TCATGCATCAATATCGTGAT |
| *OrcuKAI2d7* | CACCATGAACATAGTTGGAGCA | TCAGGCGTCAATGATGTC |
| *OrcuKAI2d8* | CACCATGAGCACAGTTGGAGC | TCAGGCTATATCGTGTTGTAT |

**Table S5**. Sizes of exons and introns for *KAI2* genes from *O. cernua* and *O. cumana*. Missing data is indicated by -. * indicates truncated gene.

| Gene | Exon 1 | Intron | Exon 2 | Coding region | Genomic |
|---|---|---|---|---|---|
| | Number of nucleotides | | | | |
| *OrceKAI2c* | 374 | - | 442 | 816 | - |
| *OrceKAI2d1* | 373 | 158 | 441 | 814 | 972 |
| *OrceKAI2d2* | 377 | 319 | 448 | 825 | 1144 |
| *OrceKAI2d3* | 372 | - | 444 | 816 | - |
| *OrceKAI2d4* | 371 | 261 | 448 | 819 | 1080 |
| *OrceKAI2d5* | 369 | 158 | 447 | 816 | 980 |
| *OrceKAI2d6* | 324 | * | * | 324 | 551 |
| *OrcuKAI2c* | 374 | 502 | 442 | 816 | 1318 |
| *OrcuKAI2d1* | 372 | 123 | 447 | 819 | 942 |
| *OrcuKAI2d2* | 357 | 660 | 447 | 804 | 1464 |
| *OrcuKAI2d3* | 372 | 256 | 447 | 819 | 1075 |
| *OrcuKAI2d4* | 371 | 319 | 448 | 819 | 1138 |
| *OrcuKAI2d5* | 379 | 475 | 446 | 825 | 1300 |
| *OrcuKAI2d6* | 372 | 289 | 444 | 816 | 1105 |
| *OrcuKAI2d7* | 369 | 169 | 447 | 816 | 980 |
| *OrcuKAI2d8* | 369 | >1000 | 441 | 810 | >1810 |

**Table S6**. Posterior summaries of model parameters predicting seed germination in response to either DCL or Oro stimulants in *O. cernua* x *O. cumana* segregants.

| **DCL Response** | post_mean* | 2.50% | 97.50% |
|---|---|---|---|
| Intercept | -1.649 | -2.456 | -0.877 |
| *OrceKAI2d1* | 0.599 | 0.100 | 1.089 |
| *OrceKAI2d2* | 0.024 | -0.453 | 0.485 |
| *OrceKAI2d3* | -0.496 | -0.981 | -0.008 |
| *OrceKAI2d4* | -0.702 | -1.040 | -0.365 |
| *OrceKAI2d5* | 0.148 | -0.236 | 0.527 |
| *OrceKAI2d6* | -0.697 | -1.062 | -0.331 |
| *OrcuKAI2d1* | -0.670 | -1.161 | -0.191 |
| *OrcuKAI2d2* | 0.119 | -0.359 | 0.599 |
| *OrcuKAI2d3* or *OrcuKAI2d5* | 0.870 | 0.143 | 1.589 |
| *OrcuKAI2d4* | 0.362 | -0.023 | 0.751 |
| *OrcuKAI2d6* | 0.551 | -0.157 | 1.272 |
| *OrcuKAI2d7* | -0.377 | -0.827 | 0.060 |
| *OrcuKAI2d8* | 0.568 | 0.113 | 1.035 |
| Petri Dish Effect | 0.611 | 0.527 | 0.710 |
| Plant Effect | 0.736 | 0.579 | 0.922 |

| **Oro Response** | post_mean | 2.50% | 97.50% |
|---|---|---|---|
| Intercept | -0.576 | -1.317 | 0.131 |
| *OrceKAI2d1* | 0.535 | 0.032 | 1.020 |
| *OrceKAI2d2* | 0.928 | 0.423 | 1.457 |
| *OrceKAI2d3* | 0.079 | -0.364 | 0.530 |
| *OrceKAI2d4* | -0.247 | -0.640 | 0.146 |
| *OrceKAI2d5* | 0.047 | -0.330 | 0.434 |
| *OrceKAI2d6* | 0.219 | -0.151 | 0.594 |
| *OrcuKAI2d1* | -0.755 | -1.279 | -0.226 |
| *OrcuKAI2d2* | -0.207 | -0.689 | 0.273 |
| *OrcuKAI2d3* or *OrcuKAI2d5* | -0.150 | -0.860 | 0.516 |
| *OrcuKAI2d4* | 0.328 | -0.055 | 0.706 |
| *OrcuKAI2d6* | -0.156 | -0.846 | 0.521 |
| *OrcuKAI2d7* | -0.478 | -0.898 | -0.065 |
| *OrcuKAI2d8* | -0.375 | -0.803 | 0.058 |
| Petri Dish Effect | 0.569 | 0.483 | 0.672 |
| Plant Effect | 0.777 | 0.594 | 0.988 |

*post_mean indicates the mean of the posterior distribution on a particular effect, and “2.5%” (“97.5%”) indicates, the lower (upper) bound of a 95% symmetric posterior credible interval. Intervals not containing zero may be deemed as significant. Larger magnitudes indicate a larger effect size.

```
CLUSTAL format alignment by MAFFT L-INS-i (v7.429)

O. cernua     MAVATTTLTDLPDVIVSNIIAAVCDVRSRNSAALVCRKWYVLERATRSSLCLRGNLRDLF
O. cumana     MAVATTTLTDLPDVIVSNIIAAVCDVRSRNSAALVCRKWYVLERATRSSLCLRGNLRDLF

O. cernua     MLPTCFQSVSHLDLSLLSPYGHPLTSASDPDPALIAHLLRHALPSVTSLTLYARNPSTIQ
O. cumana     MLPTCFQSVSHLDLSLLSPYGHPLTSASDPDPALIAHLLRHALPSVTSLTLYARNPSTIQ

O. cernua     LIAPQWPNLEHLKLVRWHQRPQTDDAGDELKILISECGQLKSLDLSAFYCWTDDVPLALE
O. cumana     LIAPQWPNLEHLKLVRWHQRPQTDDAGDELKILISECGQLKSLDLSAFYCWTDDVPLALE

O. cernua     FCPTFASILTCLNLLNSSFSEGFKSDEVKVITKACPNLREFRAACMFDPRYIGCVGDEAL
O. cumana     FCPTFASILTCLNLLNSSFSEGFKSDEVKVITKACPNLREFRAACMFDPRYIGCVGDEAL

O. cernua     VSVSVNCPKLAILHLADTSALSSARGDFDMEHQVLTQEDARINAATLIEVFSGLPRLEEL
O. cumana     VSVSVNCPKLAILHLADTSALSSARGDFDMEHQVLTQEDARINAATLIEVFSGLPRLEEL

O. cernua     AIDVSVNVRDSGPALEVLKSKCPGLRSLKLGQFHGISLPVGSKLDGVALCHGLKSLSIRN
O. cumana     AIDVSVNVRDSGPALEVLKSKCPGLRSLKLGQFHGISSPVGSKLDGVALCHGLKSLSIRN

O. cernua     VSDLSDMGLIAIGRGCCRLAKFEVHGCRKLTVRGLRTMASLLHRTLVDVRISCCKSLGAV
O. cumana     VSDLSDMGLIAIGRGCCRLAKFEVHGCRKLTVRGLRTMASLLHRTLVDVRISCCKSLGAV

O. cernua     QSLQALEPLQDRIERLHIDCIWDCTTDELDETNDDDCFDLKSSDQGGVLNSYQPDEHTAQ
O. cumana     QSLQALEPLQDRIERLHIDCIWDCTTDELDETNDDDCFDLKSSDQGGVLNSYQPDEHTAQ

O. cernua     EWTGTDYDYDYDGMTHAIKKRKCSHDQNPSYFGMVVNSNGSENVNAYGERVWDRLQCLSL
O. cumana     EWTGTDYDYDYDGMTHAIKKRKCSHDQNPSYFGMVVNSNGSENVNAYGERVWDRLQCLSL

O. cernua     SVPVGQLLNPLVSAGLENCPNLEEIRIKIEGDCRVLPKPTVREFGLSTLVIYPSLSKMHL
O. cumana     WVPVGQLLNPLVSAGLENCPNLEEIRIKIEGDCRVLPKPTVREFGLSTLVIYPSLSKMHL

O. cernua     DCGDIIGYTHTAPSGQMDLSLWERFCLIGIGNLSLTELDYWPPQDRDVNQRTLSLPAAGL
O. cumana     DCGDIIGYTHTAPSGQMDLSLWERFCLIGIGNLSLTELDYWPPQDRDVNQRTLSLPAAGL

O. cernua     LQQCFGLRKLFIHGTAHEHFMMFLLRIPDLRDVQLREDYYPAPENDMSTEMRADSCSRFE
O. cumana     LQQCFGLRKLFIHGTAHEHFMMFLLRIPDLRDVQLREDYYPAPENDMSTEMRADSCSRFE

O. cernua     VALNGRQISD
O. cumana     VALNGRQISD
```

**Figure S1**. Alignment of MAX2 protein sequences from *O. cernua* and *O. cumana*. CLUSTAL format alignment by MAFFT, with disagreements highlighted.

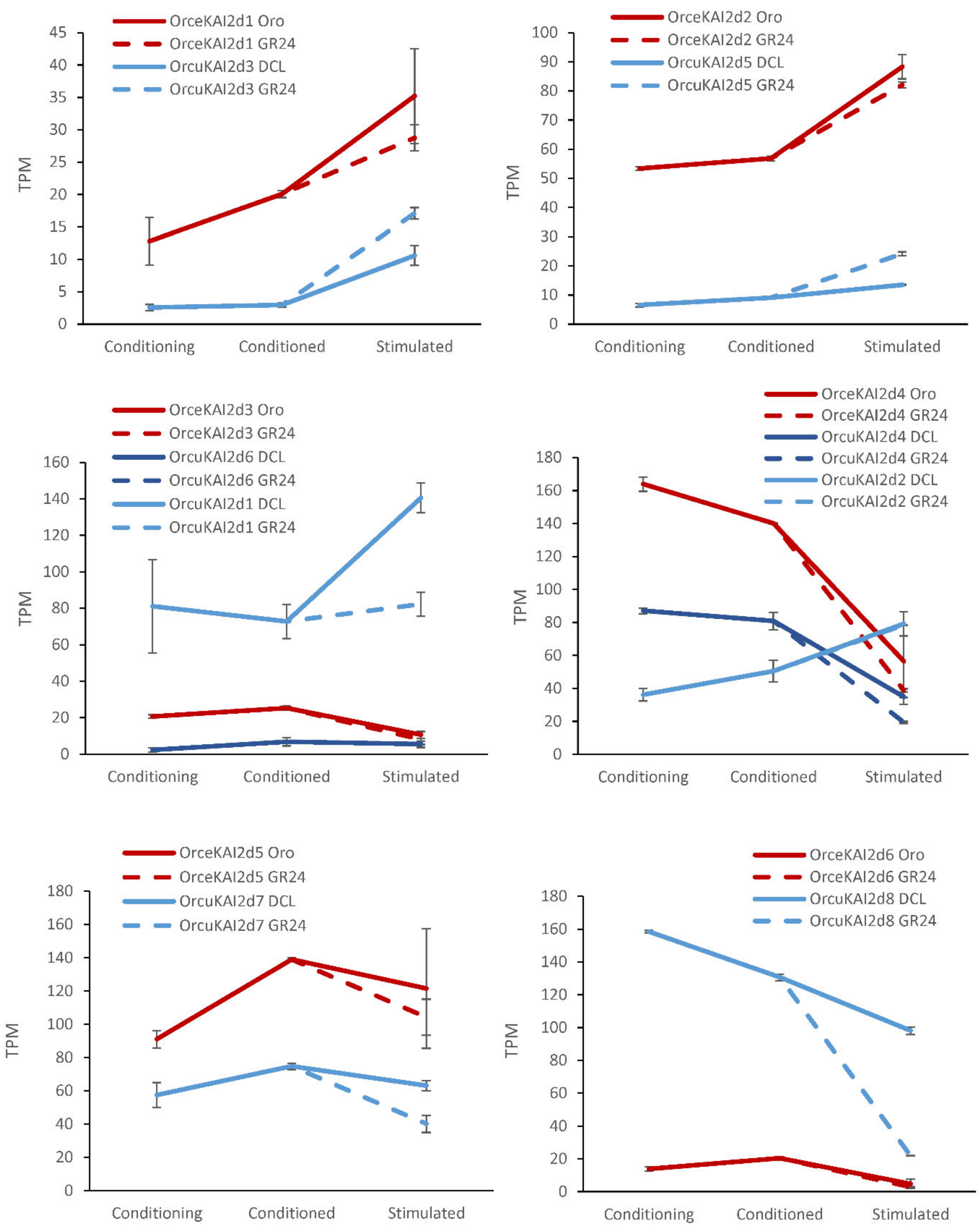


**Figure S2.** *KAI2* gene expression from each transcriptome stage. For each species, solid line indicates expression in response to species-specific stimulant (orobanchol or DCL) and dotted line indicates expression in response to universal stimulant GR24. For ease of visualization, the genes are distributed among multiple graphs and grouped according to putative othologs for *O. cernua* (Orce) and *O. cumana* (Orcu) as depicted in Fig. 1.

### a) Response to Orobanchol

*O. cernua* genes

*O. cumana* genes

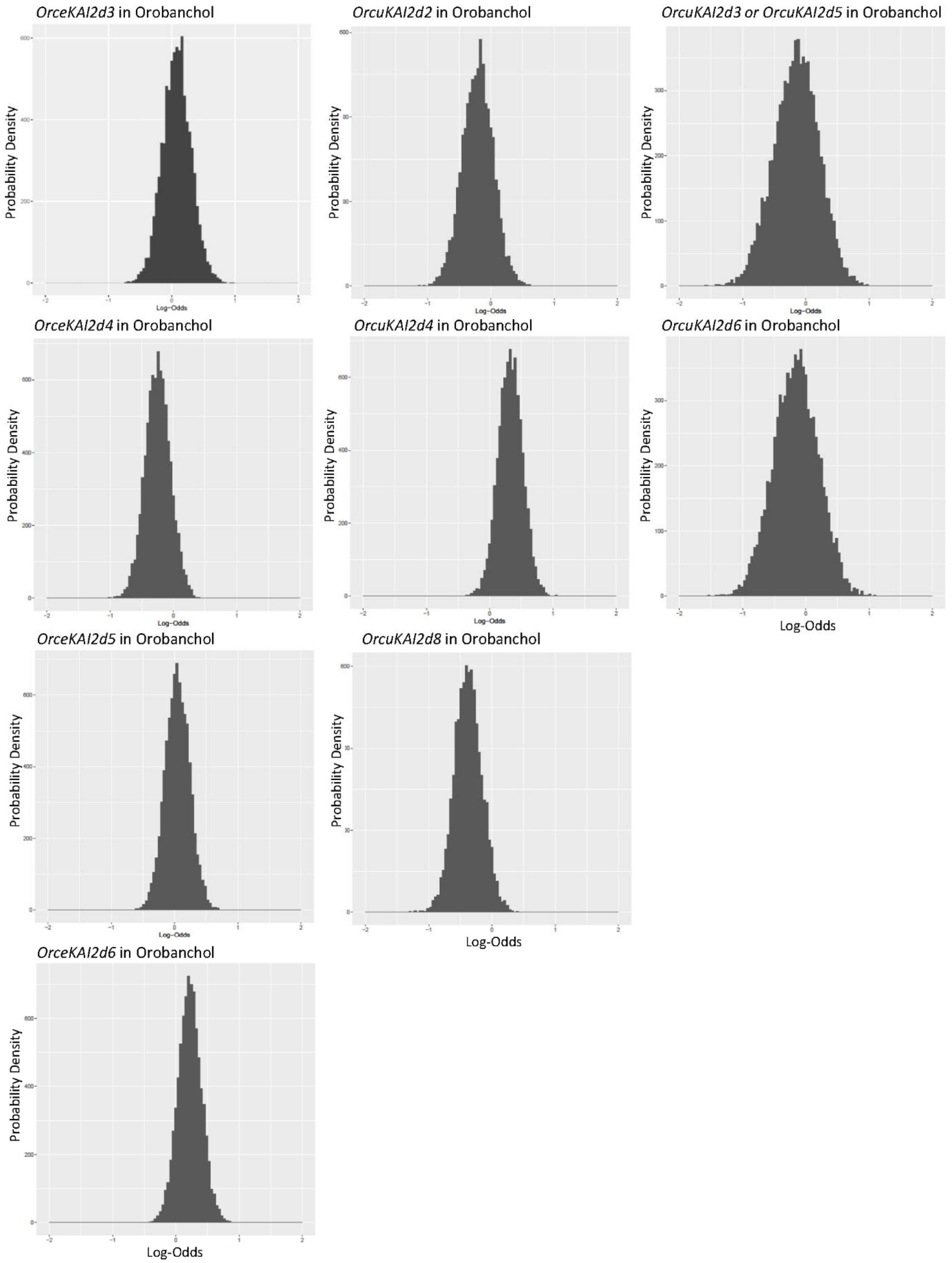

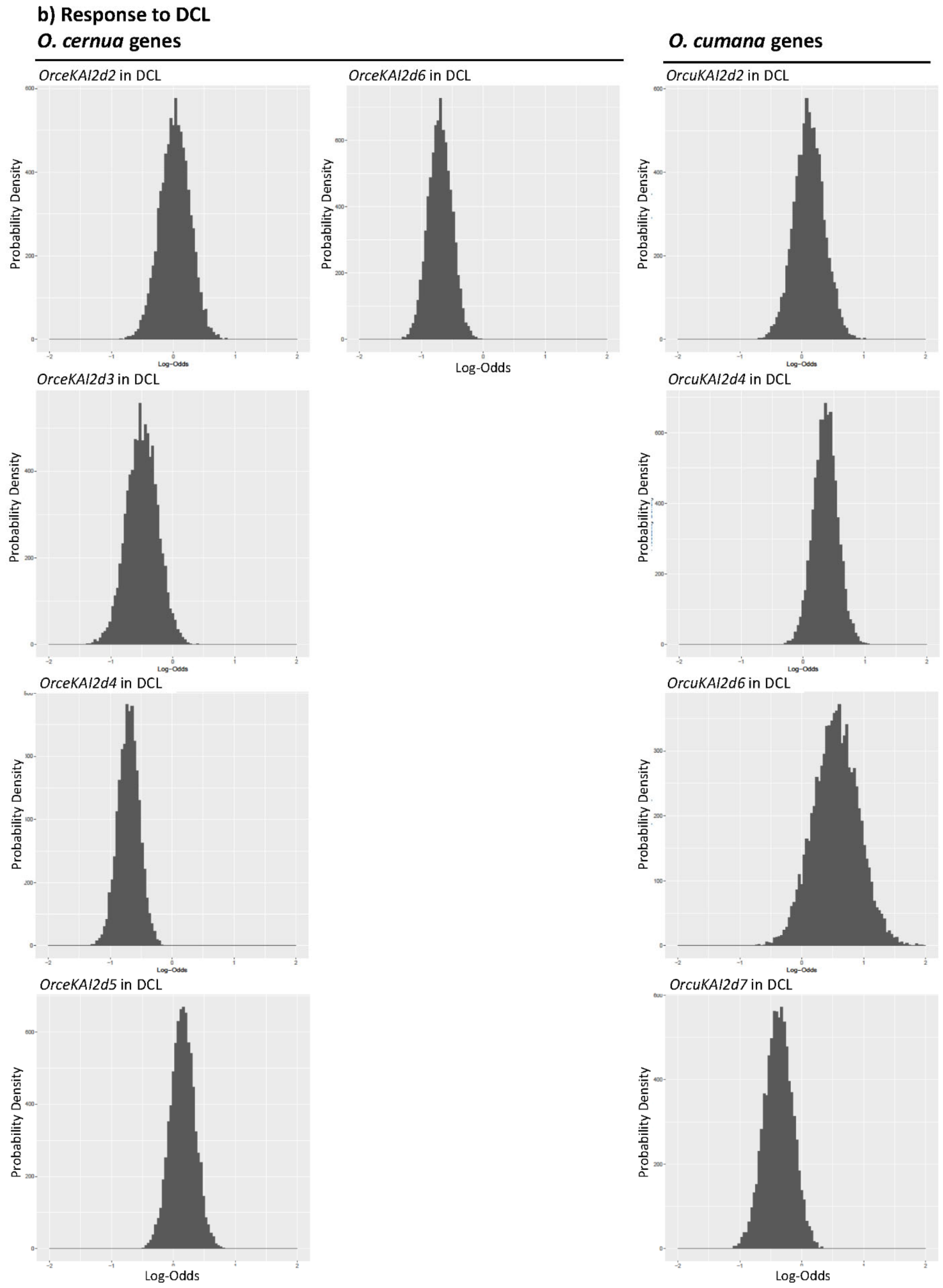


**Figure S3**. Posterior distribution charts for (a) *KAI2d* genes in response to orobanchol and (b) *KAI2d* genes in response to DCL. Higher values of density indicate that it is more probable that the truth lies in that region. Sign of log-odds indicates direction of correlation, and magnitude indicates strength of correlation. Values far removed from 0 indicates correlation of gene with germination response to a given phenotype.

CLUSTAL format alignment by MAFFT (v7.475)

```
               1          10         20         30         40         50
               |          |          |          |          |          |
Arabid_D14     M-SQHNILEALNVRVVGTGDRILFLAHGFGTDQSAWHLILPYFTQNYRVVLYDLVCAGSV
Arabid_KAI2    M-GVV--EEAHNVKVIGSGEATIVLGHGFGTDQSVWKHLVPHLVDDYRVVLYDNMGAGTT
OrceKAI2c      M-GIT--QDAHNVRFLGSGSRTIVLAHGFGTDQSVWKHLVPHLVGEYRVVLYDNMGAGTT
OrcuKAI2c      M-GIT--Q?AHNVR?LGSG?RTIVLAHGFGTDQSVWKHLVPHLVGEYRVVLYDNMGAGTT
OrceKAI2d1     MSSIV--GARHNARVLGSGQTTVVLGHGFGTDQSVWRHLVPRLVDQYRVVLYDNMGAGTT
OrceKAI2d2     MNSIVGLGAAHNVRVLGSGQTTVVLNHGFGTDQSVWRHLVPHLVDQYRVVLFDNMGAGTT
OrceKAI2d3     MSSIV--GARHNVRVLGSGQTTVVL?HGFGTDQSVWRHLVPRLVDQYRVVLYDNMGAGAT
OrceKAI2d4     MSSIV--GAAHNVRVLGSGQTTVVLGHGYGMDQSVWGHLVPRLVDHYRVVLYDNMGAGTT
OrceKAI2d5     M-NIV--GAAHNVRVLGSGEATVVLGHVFGTDQSIWRHLVPHLVDHYRVVLYDNMGAGTT
OrceKAI2d6     M-STV--GADHNGRVLGSGETTIVLGHGYGTDQSVWRYLVPHLVDLYRVILYDNMGAGTT
OrcuKAI2d1     MSSIV--GARHNVRVVGSGQTTVVLGHGFGVDQSVWRHLAPRLVDQYRVVLYDNMGAGTT
OrcuKAI2d2     MGSIV--GAAHNVRV-----TTVVLGHGLGIDQSVWRHLVPRLVDHYKVVLYDNMGAGTT
OrcuKAI2d3     MSSIV--GARHNVRVLGSGQTTVVLGHGFGTDQSVWRHLVPRLVDQYRVVLYDNMGAGTT
OrcuKAI2d4     MSSIV--GAAHNVRVLGSGQTTVVLGHGYGMDQSVWRHLVPRLVDHYRVVLYDNMAAGTT
OrcuKAI2d5     MNRIVGLGAAHNVRVLGSGQTTVVLNHGFGTDQSVWRHLVPHLVDQYRVVLFDNMGAGTT
OrcuKAI2d6     MSSIV--GARHNVRVLGSGPTTVVLAHGFGTDQSVWRHLVPRLVDQYRVVLYDNMGAGTT
OrcuKAI2d7     M-NIV--GAAHNVRVLGSGETTVVLGHGFGTDQSIWRHLVPHLVDQYRVVLYDNMGAGTT
OrcuKAI2d8     M-STV--GADHNVRVLGSGETTIVLGHGYGTDQSVWRYLVPHLVDLYRVILYDNMGAGTT

               60         70         80         90         100        110
               |          |          |          |          |          |
Arabid_D14     NPDYFDFNRYTTLDPYVDDLLNIVDSLGIQN-CAYVGHSVSAMIGIIASIRRPELFSKLI
Arabid_KAI2    NPDYFDFDRYSNLEGYSFDLIAILEDLKIES-CIFVGHSVSAMIGVLASLNRPDLFSKIV
OrceKAI2c      NPDYFDFERYSTLEGFAYDVIAILEELEISSGCIYVGHSVSAMIGIIASLTRPDLFSKIL
OrcuKAI2c      NPDYFDFERYSTLEGFAYDVIAILEELEISSGCIYVGHSVSAMIGIIASLTRPDLFSKIL
OrceKAI2d1     NPNNYDFDRYATLDGHANDLLAILEEFSIGK-CIHVGHSLSAMVGAMASISRPNLFHKLV
OrceKAI2d2     NPDYYDFERYATLEGYADDLLAILEEFSIGK-CIYVGHSLSAMVGALASIFRPDLFHKLI
OrceKAI2d3     NPDSYDFDRYAILDGHANDLLAILEEFSV?K-CIHVGHSLSAMVGAMASILRPDLFHKLV
OrceKAI2d4     NPNNYDFDRYATLDGFANDLISILEEFSLGK-CIYVGHSLSAMVGALASVLRPNLFHKLV
OrceKAI2d5     NPDLYDFDRYSTLEGHAYDLIAILEELSNGK-CIHVGHSLSAMVAATASIFRPDLFHKLV
OrceKAI2d6     NPD-----RYGPLCEHRRV-------------CLRFARHFGGIRERKMHICRP-----LI
OrcuKAI2d1     NPDSYDFDRYATLDGHASDLLAILEEFSVGK-CIHVGHSLSAMVAAMASISRPDLFHKLI
OrcuKAI2d2     NPDKYDFNRYATLNGYADDLLAILEEFSVEK-CIYVGHSFSAMVGAMASILRPDLFRKLI
OrcuKAI2d3     NPDSYDFDRYAVLDGHANDLLAILEEFSIGK-CIHVGHSMSAMVGAMASISRPNLFHKLV
OrcuKAI2d4     NPNNYDFDRYATLDGFANDLISILEEFSIGK-CIYVGHSLSAMVGALASVLRPNLFHKLV
OrcuKAI2d5     NPDYYDFERYAALEGYADDLLAILEEFSIEK-CIYVGHSMSAMVGAMASIFRPDLFHKLI
OrcuKAI2d6     NPDSYDFDRYAILDGHANDLLAILEEFSVEK-CIHVGHSLSAMVGAMASILRPDLFHKLV
OrcuKAI2d7     NPDLYDFDRYSTLEGHAYDLIAILEEFSNGK-CIHVGHSLSAMVAATASIFRPDLFHKLV
OrcuKAI2d8     NPDRYDFDRYASIEGHASDLLAILEEFGSGK-CIFVGHSLSCMAAALASIYRPDLFHKLV

                120        130        140        150        160        170
                |          |          |          |          |          |
Arabid_D14     LIGFSPRFLNDEDYHGGFEEGEIEKVFSAMEANYEAWVHGFAPLAVGADVPA-AVREFSR
Arabid_KAI2    MISASPRYVNDVDYQGGFEQEDLNQLFEAIRSNYKAWCLGFAPLAVGGDMDSIAVQEFSR
OrceKAI2c      TISGSPRYLNDPDYVGGFGQDELRELFDAMKSNYRAWCLGFAPLCVGGDMESAAVQEFSR
OrcuKAI2c      TISGSPRYLNDPDYVGGFGQDELRELFDAMKSNYRAWCLGFAPLCVGGDMESAAVQEFSR
OrceKAI2d1     MISATPRTVSTEDYYGGLNQEDLDQLLDAMETNHNSMLHGLAPLAIGGDMDSEVVQEYSR
OrceKAI2d2     MISASPRMLNTEGYYGGLEQKDHDQLLVAMETNFKSLVEGSAPLVIGGDMDSEVVQEYSR
OrceKAI2d3     MISATPRVVNTEDYYGGLNQEDLDQILEAMETNYNSMVHGMAPLAIGGDMDSEVVQEYSR
OrceKAI2d4     MLSAIPRMSNTEDYYGGFNQEDIDQLIAGMETNYNSMIHGMAPLVIGGDMDSEAVQEYSR
OrceKAI2d5     MMSATPRTSNTADYYGGMEQKDMDQILDAMETNLESLISGWAPLAVGGDMDSPTVQEFSR
OrceKAI2d6     VLHGS----------------------------------------SLGFDIPT*SVS*AGH
OrcuKAI2d1     MISTTPRVVNTEDYYGGFNQEDLDQLVEAMETNYNSMLHGVAPLMIGGDMDSEVVQEYSR
OrcuKAI2d2     MIAATPRMSNTEDYYGGLNQEDVDQLLYGLETNHNSMLHGLAPLVIGGDMDSEVVQEYSR
OrcuKAI2d3     MISATPRMASTEDYYGGLNQEDLDQLLDAMETNHNSMLHGLAPLVIGGDMDSEVVQEYSR
OrcuKAI2d4     MLSAIPRMSNTEDYYGGFNQEDIDQLIAGMETNYNSMIHGMAPLVIGGDMDSEAVQEYSR
OrcuKAI2d5     MMSATPRSTNTEGYYGGLDQKDLDQVLVAMETNYKSVMEGLAPLVIGGDMDSEAIQEFSR
OrcuKAI2d6     MISATPRVVNTEDYYGGLNQEDLDQILEAMETNYNSMVHGMATLAIGGDMDSEVVQEYSR
OrcuKAI2d7     MMSATPRASNTADYYGGMEQKDIDQILDAMETNIESLISGWAPLAVGGDMDSPAVQEFSR
OrcuKAI2d8     MLCATPRMSNTVDYYGGFPQEEINQLLDAMATNYESYTLGMAPLAIGCDLDSEALQEYSR
```

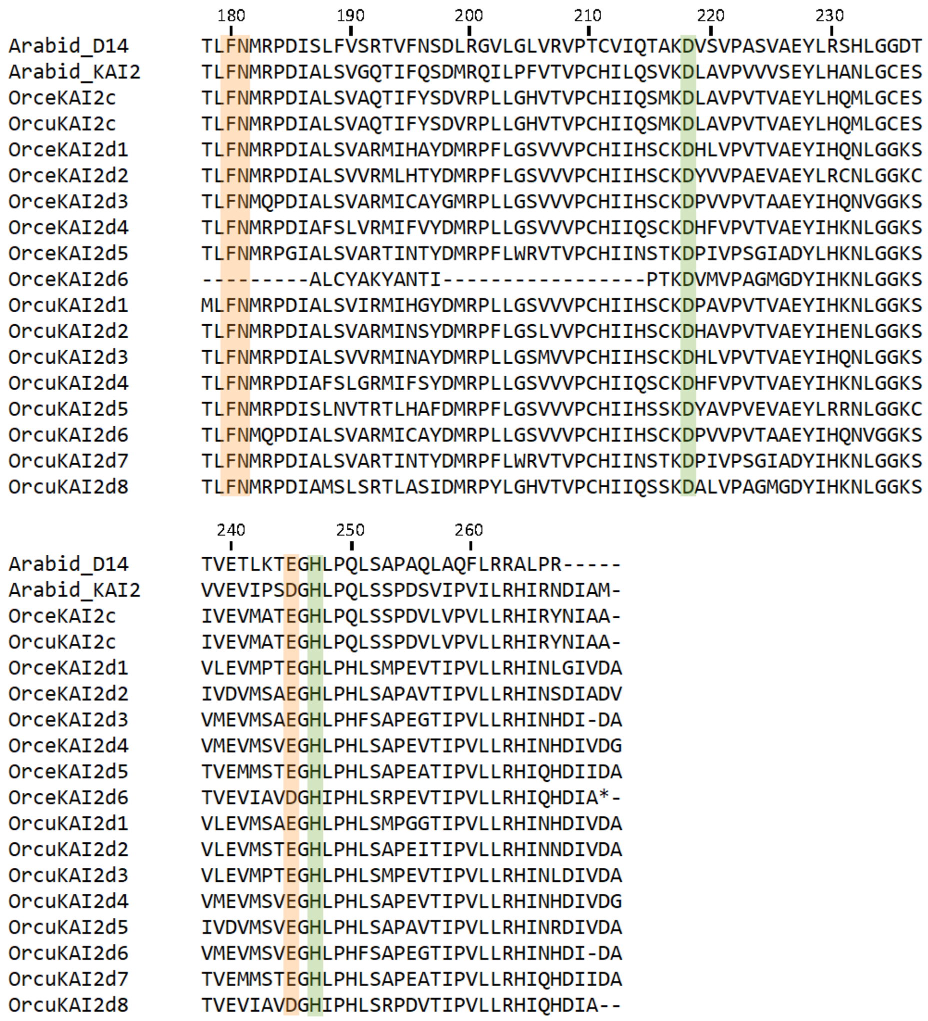


**Figure S4.** Alignment of all KAI2 protein sequences from *O. cernua* and *O. cumana*, along with KAI2 and D14 from *Arabidopsis thaliana*. Conserved amino acids indicated as being essential for SL recognition (as determined in *A. thaliana*) are highlighted in green. Amino acids indicated as important for association with D3 (*Oryza sativa* homolog of *A. thaliana* MAX2) (Yao et al., 2016) are highlighted in orange. Numbering is according to *A. thaliana* D14.

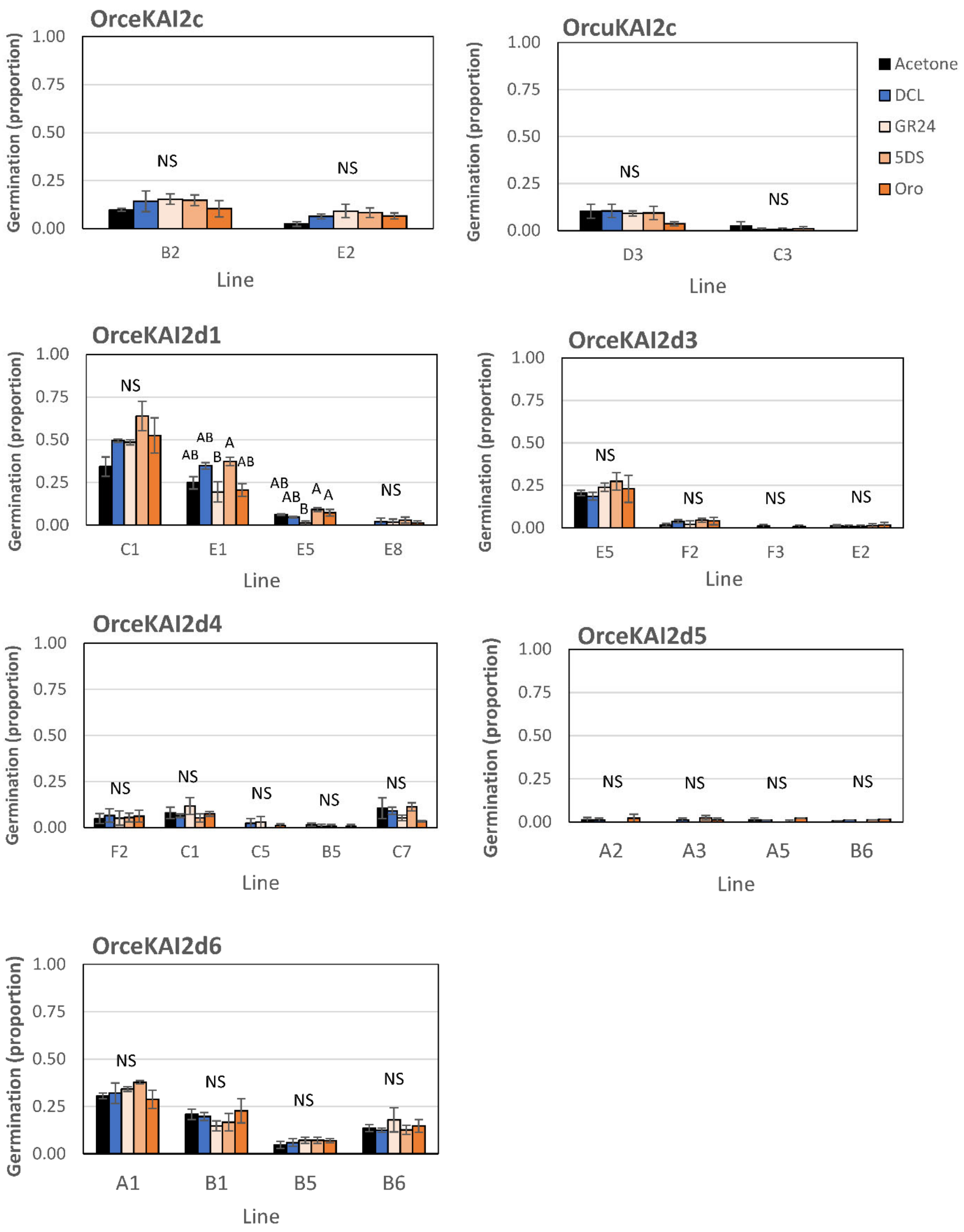
OrceKAI2c
Germination (proportion)
1.00
0.75
0.50
0.25
0.00
NS
NS
B2
E2
Line
OrcuKAI2c
Germination (proportion)
NS
NS
D3
C3
Line
Acetone
DCL
GR24
5DS
Oro
OrceKAI2d1
Germination (proportion)
NS
AB
AB
B
A
AB
AB
AB
B
A
A
NS
C1
E1
E5
E8
Line
OrceKAI2d3
Germination (proportion)
NS
NS
NS
NS
E5
F2
F3
E2
Line
OrceKAI2d4
Germination (proportion)
NS
NS
NS
NS
NS
F2
C1
C5
B5
C7
Line
OrceKAI2d5
Germination (proportion)
NS
NS
NS
NS
A2
A3
A5
B6
Line
OrceKAI2d6
Germination (proportion)
NS
NS
NS
NS
A1
B1
B5
B6
Line

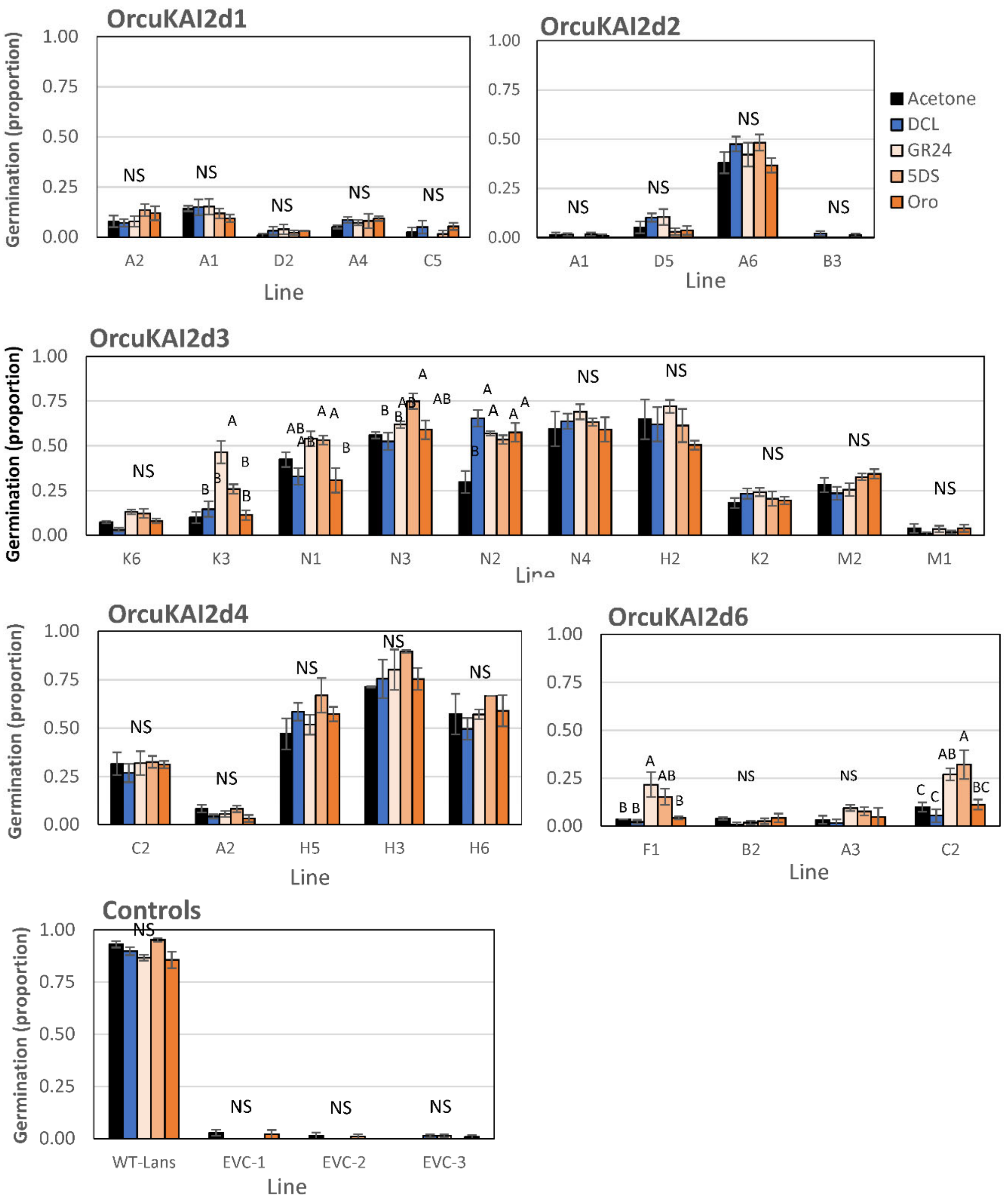


**Figure S5**. Assays of *Orobanche cernua* and *O. cumana KAI2c* and *KAI2d* gene responses to germination stimulants in a heterologous system. The indicated genes were expressed as transgenes in the *A. thaliana kai2* mutant background. Seeds were scored for germination following exposure to acetone (negative control), DCL, GR24, 5DS or Oro. Each line represents a unique transformation event. Tukey-Kramer HSD test was used to determine significance, * $P < 0.05$. Each column represents the mean of 3 replications and vertical lines represent SE.
